# The role of MICOS in modulating mitochondrial dynamics and structural changes in vulnerable regions of Alzheimer’s Disease

**DOI:** 10.64898/2025.12.13.693635

**Authors:** Bryanna Shao, Bartosz Kula, Han Le, Prasanna Venkhatesh, Prasanna Katti, Andrea G. Marshall, Siddharth Chittaranjan, Suraj Thapilyal, Namdar Hari Kalpana, C. Nivedya, Adam Roszczyk, Harrison Mobley, Mason Killion, Emelina St John, Pamela Martin, Benjamin Rodrigiuez, Markis’ Hamilton, LaCara Bell, Sunjay M. Wyckoff, Laurent A. Moran, Mark Philips, David Hubert, Briar Tomeau, Jeremiah M. Afolabi, Annet Kirabo, Ismary Blanco, Samantha Reasonover, Lauren E. Drake, Ethan S. Lippmann, Chantell Evans, Monica M. Santisteban, Taneisha G. Cheairs, Sepiso Mesenga, Celestine Wanjalla, Jennifer Gaddy, Ronald McMillan, Calixto P Hernandez Perez, Htet Htet Paing, Jenny C. Schafer, Bret Mobley, Julia Berry, Amber Crabtree, Oleg Kovtun, Shawn Goodwin, Edgar Garza Lopez, Chandravanu Dash, Dao-Fu Dai, Tyne W. Miller-Fleming, Antentor Hinton, Nathan A. Smith

## Abstract

Mitochondrial contact site and cristae organizing system (MICOS) complexes are critical for maintaining the mitochondrial architecture, cristae integrity, and organelle communication in neurons. MICOS disruption has been implicated in neurodegenerative disorders, including Alzheimer’s disease (AD), yet the spatiotemporal dynamics of MICOS-associated neuronal alterations during aging remain unclear. Using three-dimensional reconstructions of hypothalamic and cortical neurons, we observed age-dependent fragmentation of mitochondrial cristae, reduced intermitochondrial connectivity, and compartment-specific changes in mitochondrial size and morphology. Notably, these structural deficits were most pronounced in neurons vulnerable to AD-related pathology, suggesting a mechanistic link between MICOS disruption and the early mitochondrial dysfunction observed in patients with AD. Our findings indicate that the loss of MICOS integrity is a progressive feature of neuronal aging, contributing to impaired bioenergetics and reduced resilience to metabolic stress and potentially facilitating neurodegenerative processes. MICOS disruption reduced neuronal firing and synaptic responsiveness, with miclxin treatment decreasing mitochondrial connectivity and inducing cristae disorganization. These changes link MICOS structural deficits directly to impaired neuronal excitability, highlighting vulnerability to AD-related neurodegeneration. These results underscore the importance of MICOS as a critical determinant of neuronal mitochondrial health and as a potential target for interventions aimed at mitigating AD-related mitochondrial dysfunction.

## 1. Introduction

Mitochondria are central to neuronal function and regulate energy metabolism, calcium homeostasis, and apoptotic signaling, all of which are essential for synaptic activity and neuronal survival (Duchen and Szabadkai, 2010; Glancy et al., 2020., Saito, He, Yan, et al. 2016; Wang et al. 2018; Yan et al. 2015; Yang et al. 2019). Neurons are particularly sensitive to mitochondrial perturbations because of their high energy demands and limited regenerative capacity, making them vulnerable to age-related metabolic dysfunction (Rangaraju et al., 2019; Li et al., 2020). Age-associated alterations in mitochondrial dynamics, including imbalances in fission and fusion (Hinton et al. 2023; Mai et al. 2010; Santos et al. 2015; Saxton and Hollenbeck 2012; Smith and Gallo 2018; Toda et al. 2016), and disruptions in mitochondrial motility have been linked to neurodegenerative disorders, especially Alzheimer’s disease (AD) (Zhu et al., 2012; Zhang et al., 2016; Sukhorukov et al., 2024., Amorim et al. 2022., Fronczek et al. 2012; Shang et al. 2022; Stahon et al. 2016).

The mitochondrial contact site and cristae organizing system (MICOS) complex plays a pivotal role in maintaining the cristae architecture, shaping inner mitochondrial membrane morphology, and mediating interactions between mitochondria and the endoplasmic reticulum (Stephan et al., 2020; Vue et al., 2023., Friedman and Nunnari 2014). Loss or dysfunction of MICOS components leads to mitochondrial fragmentation, a reduced cristae density, impaired oxidative phosphorylation, and increased oxidative stress—all key features observed in vulnerable neurons during AD pathogenesis (Yang et al., 2015; Vue et al., 2024., Dong et al. 2022; Chan 2012; Dietrich et al. 2013; Shang et al. 2022; Vue et al. 2024). MICOS-mediated regulation of the mitochondrial ultrastructure is also critical for calcium handling and metabolic coupling, processes that are essential for maintaining hypothalamic neuronal function (Katti et al., 2023; Chen et al., 2023).

Age-related mitochondrial dysfunction in these neurons can lead to impaired synaptic activity, altered excitability, and disrupted metabolic control, potentially priming the brain for neurodegenerative processes (Newton et al., 2013; Quarta et al., 2020). Given that early hypothalamic dysfunction is increasingly recognized as a contributor to AD, understanding how MICOS integrity influences the mitochondrial structure and neuronal physiology during aging is critically important (Vercruysse et al., 2018; Ardanaz et al., 2022). In this study, we examine age-dependent alterations in the MICOS-dependent mitochondrial architecture in murine hypothalamic neurons and explore the structural features that may underlie increased neuronal susceptibility to AD-related metabolic and bioenergetic stress. Moreover, MICOS-dependent mitochondrial integrity is closely linked to neuronal electrophysiological properties. Disruptions in the crista architecture can alter ATP availability, calcium buffering, and synaptic excitability, directly affecting action potential firing and neurotransmitter release (Ghosh and Greenberg, 1995; Rintoul et al., 2003). Recent evidence also suggests that mitochondrial structural deficits can influence microglial interactions and inflammatory signaling, further modulating hypothalamic neuronal function during aging (Bu et al., 2014; Bhusal et al., 2022). Understanding these relationships provides a mechanistic framework to connect the mitochondrial ultrastructure with functional vulnerability in neurons, particularly in the context of AD.

## 2. Results

### 2.1 Demographic and Clinical Characteristics of the Study Population

We divided participants into two age groups (≤35 years, n=88; >50 years, n=268) to assess differences in the baseline demographic and biochemical characteristics. The younger cohort was composed of 67% females and 33% males, while the older group included 72.4% females and 27.6% males. This difference in sex distribution was not statistically significant (p=0.41).

Several biochemical markers varied significantly between age groups. The estimated glomerular filtration rate (eGFR) was notably higher in younger individuals (111 ± 24 mL/min/1.73 m²) than in older participants (89 ± 18 mL/min/1.73 m²; p=3.85e-12), which is consistent with the expected age-related decline in renal function. Calcium levels, which were within the normal range, were lower in the younger group (2.30 ± 0.08 mmol/L) than in the older group (2.33 ± 0.09 mmol/L, p=0.001). Similarly, the chloride concentration was slightly but significantly lower in the younger group (101.14 ± 2.09 mmol/L vs. 101.87 ± 2.47 mmol/L, p=0.007), as was the sodium concentration (139.06 ± 1.79 mmol/L vs. 140.67 ± 2.12 mmol/L, p=5.39e-11). Urea nitrogen levels increased significantly with age, with values of 11.68 ± 3.17 mg/dL in the younger group and 14.31 ± 3.98 mg/dL in the older group (p=2.03e-09). The urinary albumin level was substantially higher in older participants (26.86 ± 100.87 mg/L) than in younger individuals (10.77 ± 9.76 mg/L; p=0.013), indicating increased renal stress in older adults. Notably, no statistically significant differences in creatinine, inorganic phosphate, or potassium levels were observed between the younger and older groups.

When MICOS gene expression was evaluated, MICOS10 (logFC = 0.001, p = 0.99) and MICOS13 (logFC = 0.002, p = 0.98) expression did not show significant age-related differences. However, urea nitrogen levels were positively associated with MICOS10 expression, with higher levels observed in participants in the top quartile of MICOS10 expression (logFC = 0.15, p = 0.02). In addition, potassium levels were significantly higher in individuals with elevated MICOS10 expression (logFC = -0.11, p = 0.03) and in those with higher MICOS13 expression (logFC = 0.15, p = 0.076). These findings suggest a potential link between nitrogen metabolism, the electrolyte balance, and MICOS gene expression.

### 2.2 Associations of MICOS gene expression with AD

We leveraged genotype information and transcriptome data from the Genotype Tissue Expression (GTEX) Project to investigate the association between the expression of MICOS genes (CHCHD3, CHCHD6, and OPA1) and AD in a clinical population. These paired samples were used to model the genetically regulated gene expression (GREX) of MICOS genes across individuals within the Vanderbilt University Medical Center biobank, BioVU. Because BioVU participants have deidentified electronic health records linked to their genetic information, we can use this approach to examine the clinical correlates of MICOS GREX. We examined the associations between MICOS GREX and AD among 57,526 individuals of European genetic ancestry (Figure 1A) and 10,481 individuals of African genetic ancestry (Figure 1B). Three MICOS-related genes—CHCHD3, CHCHD6, and OPA1—were evaluated in multiple brain regions.

**Figure 1.**
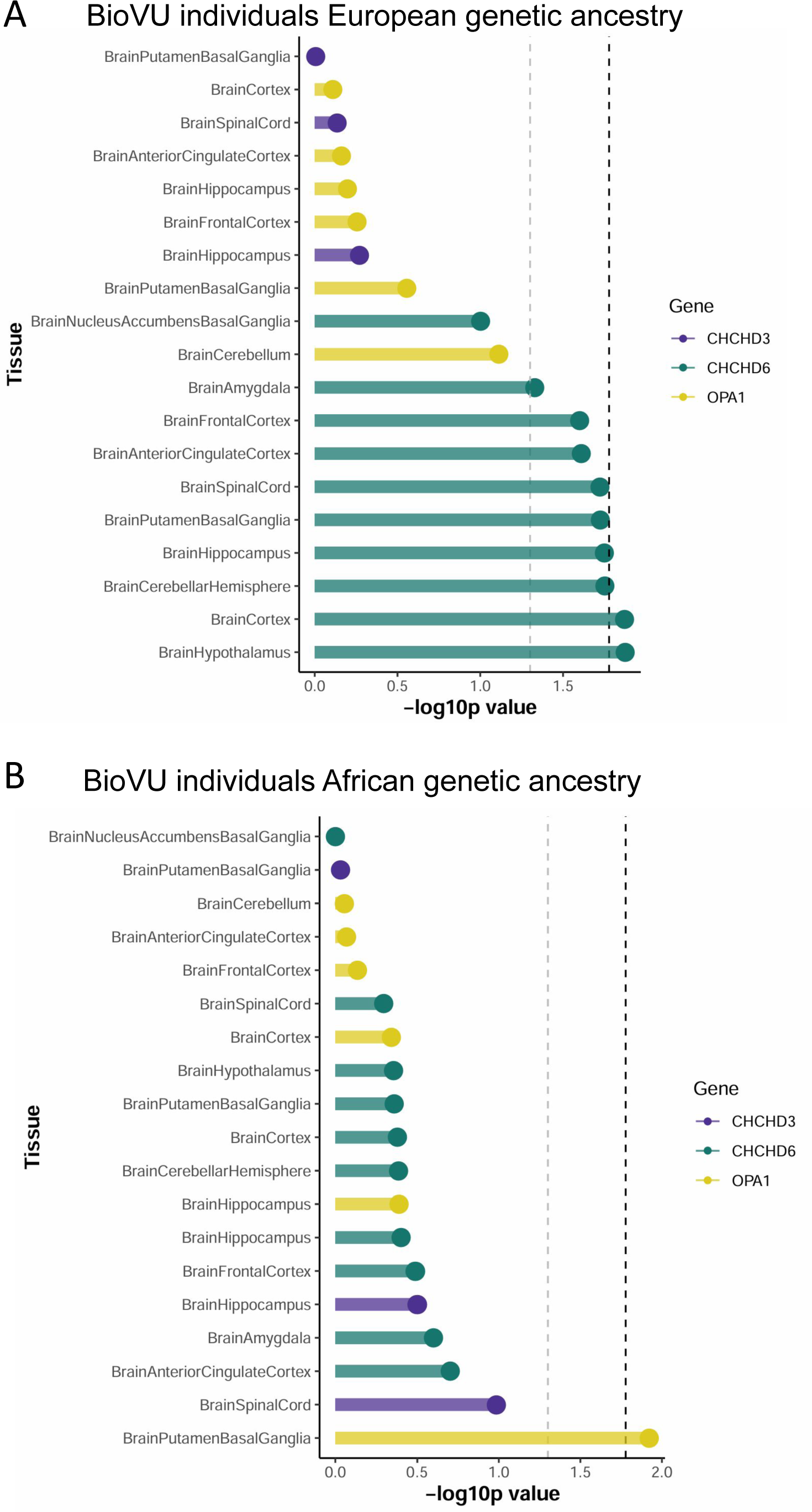
A: Associations between MICOS GREX and Alzheimer’s disease in BioVU individuals of European ancestry. GREX was calculated across 57,526 individuals of European ancestry for three genes (CHCHD3, CHCHD6, and OPA1) across all available GTEX brain tissues and tested for association with Alzheimer’s Disease. P-values are depicted with dashed lines, showing nominal values (p <0.05, gray) and the more stringent Bonferroni-corrected p-value (p < 0.0167, black). CHCHD6 GREX was significantly associated with Alzheimer’s disease status in the hypothalamus and cortex tissues. Nominal associations were observed across several brain regions, including cerebellum, hippocampus, basal ganglia, spinal cord and amygdala. **B: Associations between MICOS GREX and Alzheimer’s disease in BioVU individuals of African ancestry.** GREX was calculated across 10,481 individuals of African ancestry for three genes (CHCHD3, CHCHD6, and OPA1) across all available GTEX brain tissues and tested for association with Alzheimer’s Disease. P-values are depicted with dashed lines, showing nominal values (p <0.05, gray) and the more stringent Bonferroni-corrected p-value (p < 0.0167, black). OPA1 was significantly associated with Alzheimer’s disease in the putamen/basal ganglia tissue.

In the European ancestry population of BioVU, CHCHD6 GREX in the hypothalamus and cortex was significantly associated with AD. Nominal associations were identified in the cerebellum, hippocampus, basal ganglia, spinal cord and amygdala (Figure 1A). Among the BioVU individuals of African genetic ancestry, OPA1 GREX in the putamen/basal ganglia tissue was significantly associated with AD (Figure 1B).

### 2.3 Alterations in the MICOS complex in AD pathology

We conducted a multimodal analysis of postmortem brain tissue from individuals with AD and unaffected controls to further explore the involvement of the MICOS complex in neurodegeneration (Figure 2A–C). Immunohistochemistry revealed extensive tau accumulation in the hippocampus and hypothalamus of AD patients, in contrast to the minimal tau staining in control tissue. This marked difference supports the presence of widespread neurofibrillary pathology in AD patients (Figure 2A–C).

**Figure 2.**
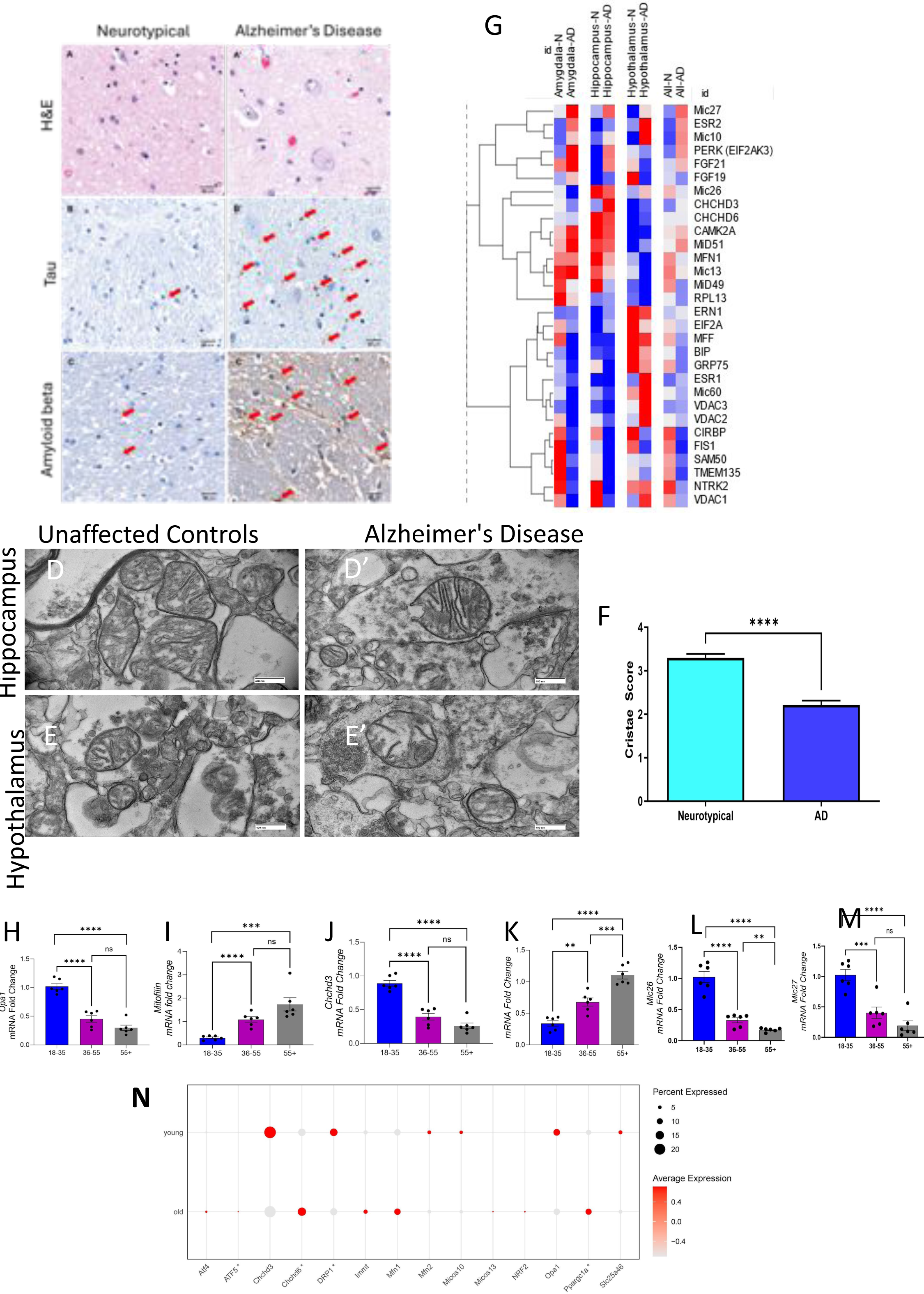
A-C: Histology staining in the hypothalamus of elderly female patients who are unaffected control or diagnosed with Alzheimer’s disease shows differences in pathology. A: Hematoxylin and Eosin staining shows cellular structure within the hypothalamus. B: Staining for tau protein in the hypothalamus. Arrows point to areas with expression of tau protein. C: Staining for amyloid beta in the hypothalamus. Arrows point to areas with expression of amyloid beta expression. D: TEM reveals cristae structure changes between neurotypical and AD patients in the hippocampus. E: TEM reveals cristae structure changes between neurotypical and AD patients in the hypothalamus. F: Cristae score quantification in the mitochondria of female neurotypical and AD patients. H: mRNA levels in OPA1, Mitofilin, CHCHD3, CHCHD6, MIC26, and MIC27 between age groups. G: Expression of MICOS complex and related mitochondrial genes in Alzheimer’s disease (AD) and controls across vulnerable brain regions. Quantification of RNA levels for numerous proteins was analyzed with a Nanodrop spectrophotometer in the amygdala, hypothalamus, hippocampus and whole brain in neurotypical and AD patients. Hierarchical clustering heat map showing differential expression of MICOS complex subunits (MIC27, MIC10, MIC26, MIC13, MIC60) and related genes regulating mitochondrial structure and dynamics (CHCHD3, CHCHD6, OPA1, MFN1, MFN2, FIS1), stress response (PERK, ERN1, GRP75, BIP, EIF2A), and mitochondrial transport/metabolism (VDAC1–3, CIRBP, TMEM135, NTRK2). The color scale indicates relative expression levels, with red representing upregulation and blue representing downregulation. F: Quantitative assessment of mitochondrial cristae integrity in postmortem brain samples from neurotypical controls and Alzheimer’s disease (AD) patients. Cristae score was significantly reduced in AD compared to neurotypical individuals (****p < 0.0001). (H–M): Quantitative RT-PCR analysis showing relative mRNA expression levels of mitochondrial dynamics and MICOS complex–related genes across three age groups (18–35, 36–55, and 55+ years). Expression levels of OPA1 (H), CHCHD3 (J), MIC26 (L), and MIC27 (M) significantly declined with advancing age, whereas CHCHD6 (I) and MIC19 (K) showed age-dependent upregulation. Data represent mean ± SEM; n = 5–6 per group; one-way ANOVA with Tukey’s post hoc test, **p < 0.01, ***p < 0.001, **p < 0.0001, ns = not significant. N: Scaled dot plot of MICOS and mitochondrial remodeling genes across young and old mouse brain samples.Dot plot showing the scaled pseudobulk expression of target genes in single-cell transcriptomes from mouse hypothalamus (Zhang et al. 2024), grouped by age class (young vs. old). Dot size indicates the percentage of cells expressing each gene, and color intensity reflects the average scaled expression level within each group. Genes significant for differential expression in pseudobulk analysis are denoted with an asterisk.

An ultrastructural analysis using electron microscopy provided direct evidence of mitochondrial abnormalities. In neurotypical brains, mitochondria displayed a well-defined morphology and intact cristae. In contrast, the mitochondria in the AD tissues exhibited marked structural disruptions, including disorganized cristae and morphological damage (Figure 2D–E). These alterations were consistently observed in both the hippocampal and hypothalamic regions. Quantitative scoring of the cristae (Figure 2F) confirmed these findings, with significantly lower scores in AD samples (2.2 ± 0.1) than in control samples (3.5 ± 0.2; p < 0.0001), underscoring the extent of mitochondrial degradation in AD patients.

### 2.4 Gene Expression

#### 2.4.1 Expression of genes involved in the MICOS complex and related mitochondrial genes in patients with AD

In both the hippocampus and hypothalamus of AD brains, strikingly similar expression patterns were observed across functional gene groups (Figure 2G). Within the MICOS complex, only the expression of MIC27 was consistently upregulated, whereas that of all the other core subunits (MIC10, MIC26, CHCHD3, CHCHD6, MIC13, MIC60, and SAM50) was substantially downregulated, suggesting a collapse of MICOS integrity despite partial compensation by MIC27. In contrast, genes regulating mitochondrial dynamics displayed a fission-dominant signature: MFN1 was downregulated, whereas FIS1, MFF, MID49, and MID51 were robustly upregulated, indicating excessive mitochondrial fragmentation. In parallel to these changes, the expression of markers of the ER stress/UPR pathway (PERK, EIF2A, ERN1, BIP, and GRP75) was uniformly upregulated, indicating an increased stress burden in both regions. Among the VDAC family members, VDAC2 was selectively upregulated, whereas VDAC1 and VDAC3 expression were suppressed, suggesting isoform-specific remodeling of outer membrane channels. Finally, in the other gene group, the expression of neurotrophic and stress-responsive regulators exhibited a mixed pattern: FGF21, FGF19, CAMK2A, TMEM135, and RPL13 were consistently upregulated, whereas CIRBP and NTRK2 were downregulated. Overall, both the hippocampus and hypothalamus exhibited a shared signature of MICOS disruption, increased mitochondrial fission, heightened ER stress, and selective remodeling of outer membrane channels, underscoring convergent mitochondrial–ER pathology in these two key brain regions of patients with AD. The trend in the amygdala was similar but less consistent and was therefore not the primary focus of this analysis.

#### 2.4.2 Age-related MICOS gene expression patterns

We examined expression patterns across three age brackets, 18–35 years, 36–55 years, and 55+ years, to assess whether the expression of MICOS genes changes with age (Figure 2H–M). The analysis of mRNA expression revealed significant group-dependent differences. *OPA1* levels were highest in the 18–35 age group and significantly decreased in both the 36–55 (p < 0.0001) and 55+ cohorts (p < 0.001), with no significant difference between the latter two groups. *Mitofilin* expression was lowest in the 18–35 age group and increased significantly in the 36–55 (p < 0.0001) and 55+ groups (p < 0.001), with no difference between the 36–55 and 55+ groups. *CHCHD3* expression was also highest in the 18–35 cohort and significantly reduced in the 36–55 (p < 0.0001) and 55+ groups (p < 0.0001), with no significant difference between the latter two groups. CHCHD6 levels were lowest in the 18–35 age group and increased significantly in both the 36–55 (p < 0.01) and 55+ cohorts (p < 0.0001), with a further significant increase between 36–55 and 55+ years (p < 0.001). *MIC26* expression was significantly decreased from 18–35 years to 36–55 years (p < 0.0001) and 55+ years (p < 0.01), with no difference between the two older groups. *MIC27* levels were significantly higher in the 18–35 age group than in both the 36–55 (p < 0.001) and 55+ cohorts (p < 0.0001), with no significant difference between the latter two groups. Together, these findings indicate that the expression of genes that regulate mitochondrial cristae follows two distinct aging trajectories: the expression of *OPA1, CHCHD3, MIC26,* and *MIC27* decreases progressively with age, whereas Mitofilin and *CHCHD6* expression increase from younger ages to midlife, with CHCHD6 expression further increasing with advanced age. This result suggests coordinated remodeling of the MICOS complex and fusion machinery across the lifespan.

#### 2.4.3 Age-Dependent Expression of MICOS and Mitochondrial Dynamics-Related Genes

We evaluated transcriptional changes in MICOS-related genes and genes related to mitochondrial dynamics with aging by analyzing single-cell RNA-seq data from the hypothalamic cells of 3-month-old (young) and 21-month-old (old) mice. The dot plot shows that the expression profiles of several MICOS complex and mitochondrial quality control genes differed between age groups (Figure 2N).

Among the MICOS genes, CHCHD6 displayed significantly higher expression in aged samples (adj. P < 0.05), whereas the changes in CHCHD3 and MIC13 expression were minimal. The expression of DRP1 and ATF5, which are associated with mitochondrial fission and stress signaling, respectively, was also significantly altered with age (adj. P < 0.05). Furthermore, the expression of the mitochondrial biogenesis regulator Ppargc1a was markedly decreased in aged cells (adj. P < 0.05), suggesting a diminished mitochondrial renewal capacity. Overall, aged hypothalamic cells exhibited a trend toward reduced expression of genes promoting mitochondrial biogenesis and fission, coupled with a modest increase in the expression of genes responsible for maintaining the mitochondrial structure, such as CHCHD6. These data indicate that transcriptional remodeling of MICOS-associated and mitochondrial dynamics pathways may contribute to an altered mitochondrial architecture and function in the aging hypothalamus.

### 2.5 Mitochondrial morphology is altered in neurons of the aging hypothalamus

Little is known about the mitochondrial structure in hypothalamic neurons. Because mitochondrial function depends on its structure, characterizing changes in the mitochondrial structure that occur during aging is crucial. We extracted and used SBF-SEM to image hypothalamic tissue from 3-month-old (n = 3) and 2-year-old male mice (n = 4), representing young (analogous to approximately 20 years in humans) and aged (analogous to approximately 70 years in humans) mice, respectively. In total, approximately 100 mitochondria were reconstructed in three dimensions for analysis, which are pseudocolored in Figure 3A–C. Using this workflow for SBF-SEM, images of orthoslices of 3-month-old and 2-year-old hypothalamic tissue were obtained (Figure 3A–A’), and 3D reconstructions of axonal mitochondria were generated (Figure 3B–C’). We arranged mitochondria based on their volume (left to right) for each age to facilitate comparisons of the mitochondrial shape across ages based on the relative size. This karyotype-like arrangement is known as “mitotyping”. Mitotyping revealed that the mitochondria of 3-month-old mice are generally rounder and less complex in shape than those of 2-year-old mice, which are elongated and contain more nanotunnels (Figure 3D–D’). The quantification of 3D reconstructed mitochondria revealed that the mitochondrial volume, area, and perimeter were lower in the hypothalamus of 2-year-old mice than in the hypothalamus of 3-month-old mice (Figure 3E–G). The 3D mitochondrial is analogous to the surface area, and the mitochondrial perimeter refers to the boundary pixel count. These alterations suggest that the size of mitochondria decreases because of aging.

**Figure 3.**
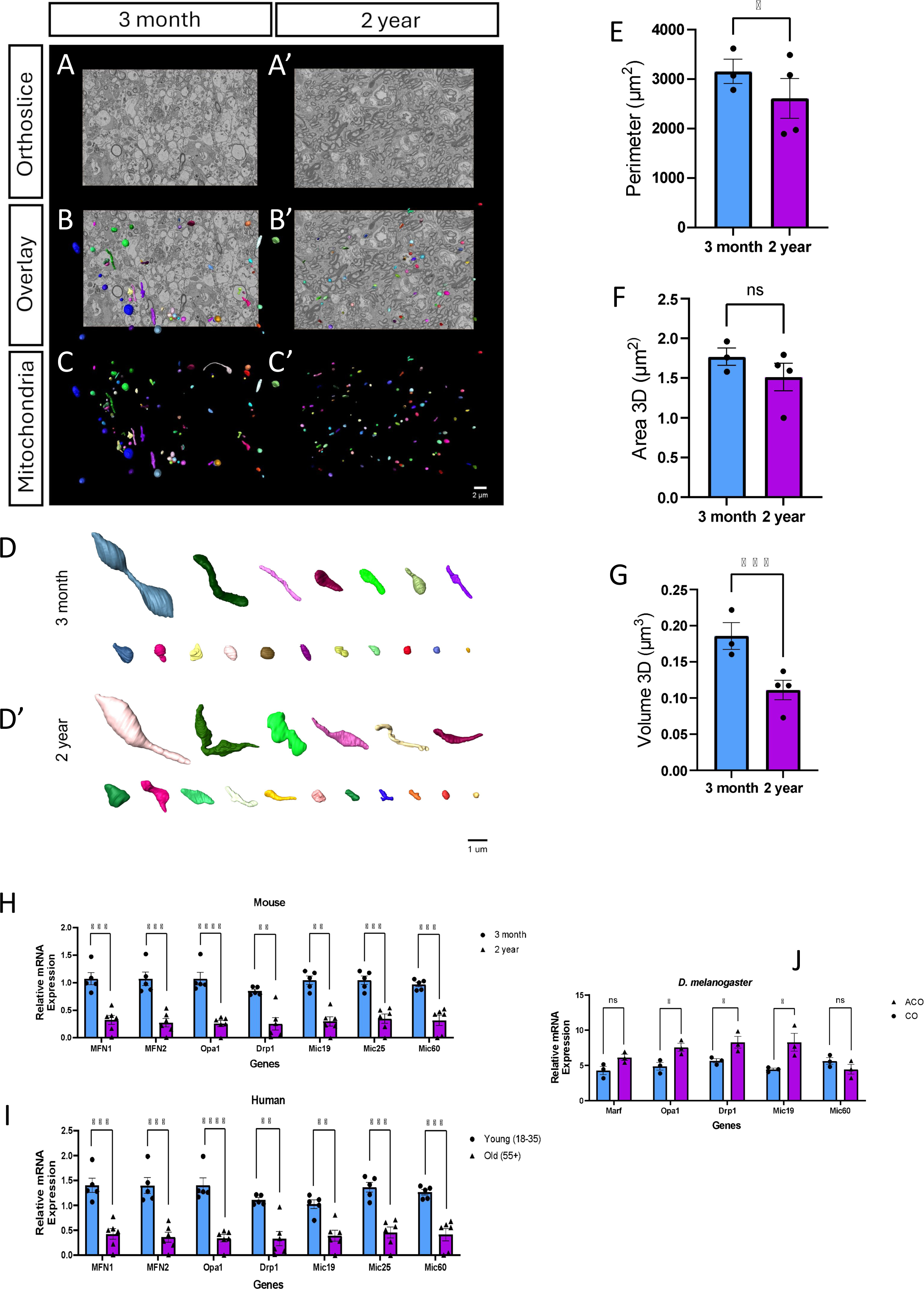
A-C: 3D morphological analysis of mitochondria within the hypothalamus of male C57Bl/6J mice for different ages. A-A’: Representative orthoslices used for manual 3D reconstruction of mitochondrial networks in the hypothalamus. B-B’: Mitochondria overlayed with an orthoslice of the hypothalamus. C-C’: Individually colored mitochondria. D: Karyotyping arranges mitochondria by relative size for mitochondria in the hypothalamus of different ages. E-G: 3D reconstructions were quantified by mitochondrial measurements in the axons of hypothalamus neurons of different ages: Perimeter (E), 3D Area (F), Volume (G). H-J: mRNA expression of mitofusin, OPA1, DRP1, MIC19, MIC25, and MIC60 in mice (H), *Drosophila melanogaster* (J), and humans (I). Young and aged cohorts were measured for mice and humans, while accelerated aging and control aging cohorts were used for *D. melanogaster.* Analysis was performed with a non-parametric Mann-Whitney test. Significance values indicate *P ≤ 0.05, ***P ≤ 0.001, and ****P ≤ 0.0001; ns, not significant.

### 2.6 Age-related decrease in the expression of the MICOS complex

Cross-species analyses are essential for determining whether mitochondrial regulatory mechanisms are conserved or represent species-specific adaptations. We addressed this question by measuring the mRNA expression of key genes regulating mitochondrial fusion (MFN and OPA1), fission (DRP1), and the MICOS complex (MIC19, Mic25, and MIC60) in *Drosophila melanogaster*, humans, and mice under comparable conditions. In *D. melanogaster* (Figure 3J), the accelerated aging cohort displayed significantly increased expression of mitochondrial remodeling genes compared with the control cohort. OPA1 expression increased by approximately 1.5–2.0-fold, DRP1 by ∼1.5-fold, and MIC19 by ∼2.0-fold (*p* < 0.05), whereas the expression of Marf (the Drosophila analog of MFN) and other MICOS components did not change significantly. These results suggest that accelerated aging in flies is associated with the upregulation of genes promoting both fusion and fission, as well as structural remodeling of cristae.

In human hypothalamic samples (Figure 3I), the direction of the effect was reversed. The expression of MFN, OPA1, and DRP1 was reduced by approximately 30–50% compared with that in the controls (*p* < 0.01). Similarly, the expression of MICOS genes (MIC19, Mic25, and MIC60) was consistently downregulated by ∼30–40% (*p* < 0.05). Mouse hypothalamic tissue displayed a highly similar pattern (Figure 3H). The expression of fusion genes (MFN and OPA1), the fission regulator DRP1, and all MICOS components was consistently downregulated under the experimental conditions, with decreases in the range of 30–50% (*p* < 0.05).

Together, these findings reveal a fundamental difference in mitochondrial gene regulation between invertebrate and mammalian systems. In *Drosophila*, accelerated aging triggers the upregulation of fusion, fission, and MICOS genes, possibly reflecting a compensatory attempt to maintain mitochondrial dynamics and cristae organization. In contrast, in both the human and mouse hypothalamus, the expression of the same gene families are significantly decreased, suggesting that aging or stress conditions suppress mitochondrial plasticity and structural maintenance in mammals. This cross-species comparison highlights the evolutionary rewiring of mitochondrial quality control and highlights the need for caution when extrapolating mechanisms from invertebrates to mammalian systems.

### 2.7 Age-Related Structural Remodeling of Axonal Mitochondria in the Hypothalamus

Axons within the SBF-SEM orthoslices were also reconstructed along with the mitochondria within them. Serial sections of mitochondria show the appearance of the nanotunnel phenotype in SBF-SEM images, along with their reconstruction within the axon (Figure 4A). Notably, we also observed nanotunnel phenotypes in mitochondria from both 3-month-old and 2-year-old mice, although more mitochondria in the 2-year-old mice formed nanotunnels. Nanotunnels are thin, double-membrane extensions in mitochondria that may play a role in direct communication between mitochondria via the exchange of matrix contents. Furthermore, we reconstructed all the mitochondria within several axons in the hypothalamus to observe the spacing within the axon in the hypothalamus of the 3-month-old and 2-year-old mice (Figure 4B and C). Mitochondria within axons in the hypothalamus tended to be fairly spaced apart with little overlap or were arranged in succession. This mirrored the compartment-specific topology of axonal mitochondria described by (Faitg et al. 2021), who used 3D EM to show that neuronal mitochondrial architecture undergoes distinct, age-dependent recalibrations across the soma, dendrites, and axons. Additionally, we did not observe any major changes in the placement or density of mitochondria within axons in the hypothalamus of aged mice. Mitochondria are known to be located throughout the axon in neurons, as well as at the growth cone at the distal tip of the axon that guides axon growth. Mitochondria are transported between the distal tip of the axon and the cell body, which contains genes that encode mitochondrial proteins and is thought to be the site of mitochondrial biogenesis and degradation. Linkage along microtubules allows mitochondria to move through anterograde and retrograde transport. Axon positioning is important for axonal homeostasis, with mitochondria being placed along parts of the axon with high metabolic demand, including synapses, growth cones, and branching sites. The mitochondrial density has also been shown to increase in axonal regions with increased activity; we observed no age-related change in the mitochondrial density, but the mitochondrial morphology changed, indicating that the structure of mitochondria rather than their number changed to accommodate changing needs in the axons of the aging hypothalamus.

**Figure 4.**
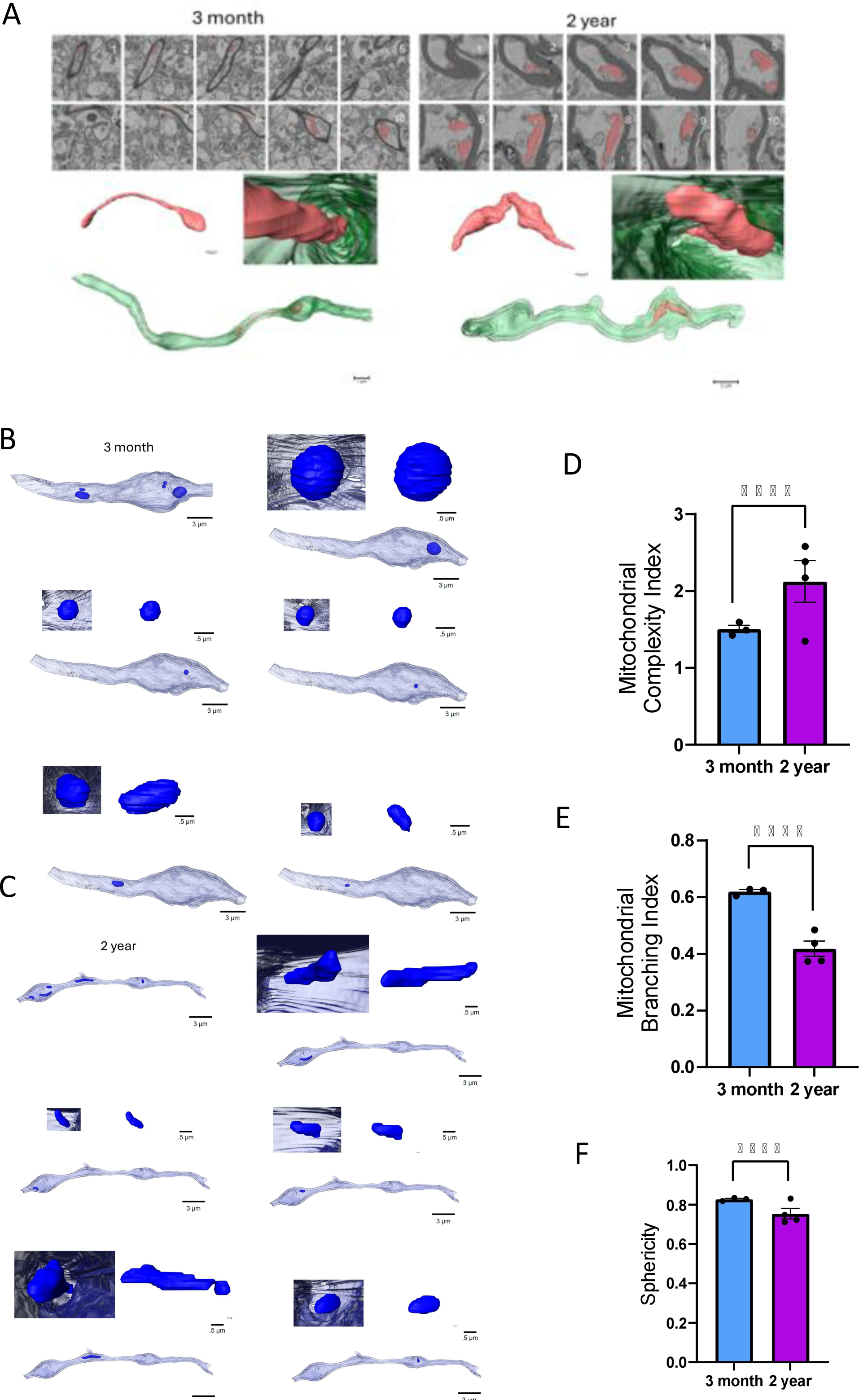
Age-Related Structural Remodeling of Axonal Mitochondria in the Hypothalamus. A: Serial sections of mitochondria with nanotunnels for 3 months and a 2-year hypothalamus on SBF-SEM images. 3D reconstructions show the mitochondria alone as well as within the axon. B-C: 3D reconstruction of an axon from a 3-month (B) and a 2-year (C) hypothalamus with each reconstructed mitochondria within the axon. D-F: Mitochondrial shape was quantified in 3D reconstructions of axonal neurons in the hypothalamus: Mitochondrial Complexity Index (D), Mitochondrial Branching Index (E), Sphericity (F). Analysis was performed with a non-parametric Mann-Whitney test. Significance values indicate *P ≤ 0.05, ***P ≤ 0.001, and ****P ≤ 0.0001; ns, not significant.

We then shifted our focus to measuring mitochondrial complexity (Figure 4D). The MCI increases with greater complexity, such as branching and an increased surface-to-volume ratio. Mitochondrial complexity can elucidate whether certain mitochondrial phenotypes are more common, which may serve specific purposes and can be indicative of certain stimuli associated with the aging process. We found that the MCI was greater in the axonal mitochondria of 2-year-old mice than in those of 3-month-old mice. The mitochondrial branching index (MBI) is the ratio of transverse to longitudinal branching, which quantifies the orientation of the branching pattern. Here, we observed that the MBI decreased in the aged hypothalamus (Figure 4E). We also considered sphericity, which is a general measurement of the roundness of mitochondria based on the ratio between volume and surface area. We found that sphericity decreased in 2-year-old mice compared with that in 3-month-old mice, which was consistent with the observed increase in elongated mitochondria (Figure 4F).

### 2.8 Electrophysiological studies

#### 2.8.1 MIC60 inhibition differentially alters the membrane properties of cortical and hypothalamic neurons

We assessed the effects of MIC60 inhibition on neuronal membrane properties by analyzing pyramidal neurons from two brain regions: the cortex, a standard model system in metabolic research, and the hypothalamus, an evolutionarily conserved structure where MIC60 disruption markedly altered mitochondrial morphology (Figure 5A–C). In cortical pyramidal neurons, MIC60 inhibition induced a significant depolarization of the resting membrane potential (RMP), increasing from −78.74 ± 1.98 mV in controls to −70.91 ± 2.46 mV following miclxin treatment (Figure 5D; Student’s *t* test, *p* = 0.017). A similar effect was not observed in hypothalamic neurons, where the RMP remained unchanged between groups (−-62.88 ± 1.56 mV vs. −60.25 ± 1.51 mV; Student’s *t* test, *p* = 0.236). Membrane resistance (R_m_) significantly increased in both regions in response to miclxin treatment. In the cortex, R_m_ increased from 239.12 ± 23.58 MΩ to 350.72 ± 40.01 MΩ (Figure 5E; Welch’s *t* test, *p* = 0.021), whereas in the hypothalamus, R_m_ increased from 868.04 ± 105.39 MΩ to 1375.76 ± 128.00 MΩ (Student’s *t* test, *p* = 0.0046). No significant differences were observed in membrane capacitance (C_m_), which remained comparable between the groups in both the cortex (129.92 ± 13.34 pF in the control vs. 130.24 ± 10.21 pF in the miclxin group; Student’s *t* test, *p* = 0.985) and the hypothalamus (47.87 ± 2.50 pF vs. 48.44 ± 2.69 pF; Student’s *t* test, *p* = 0.879; Figure 5F).

**Figure 5:**
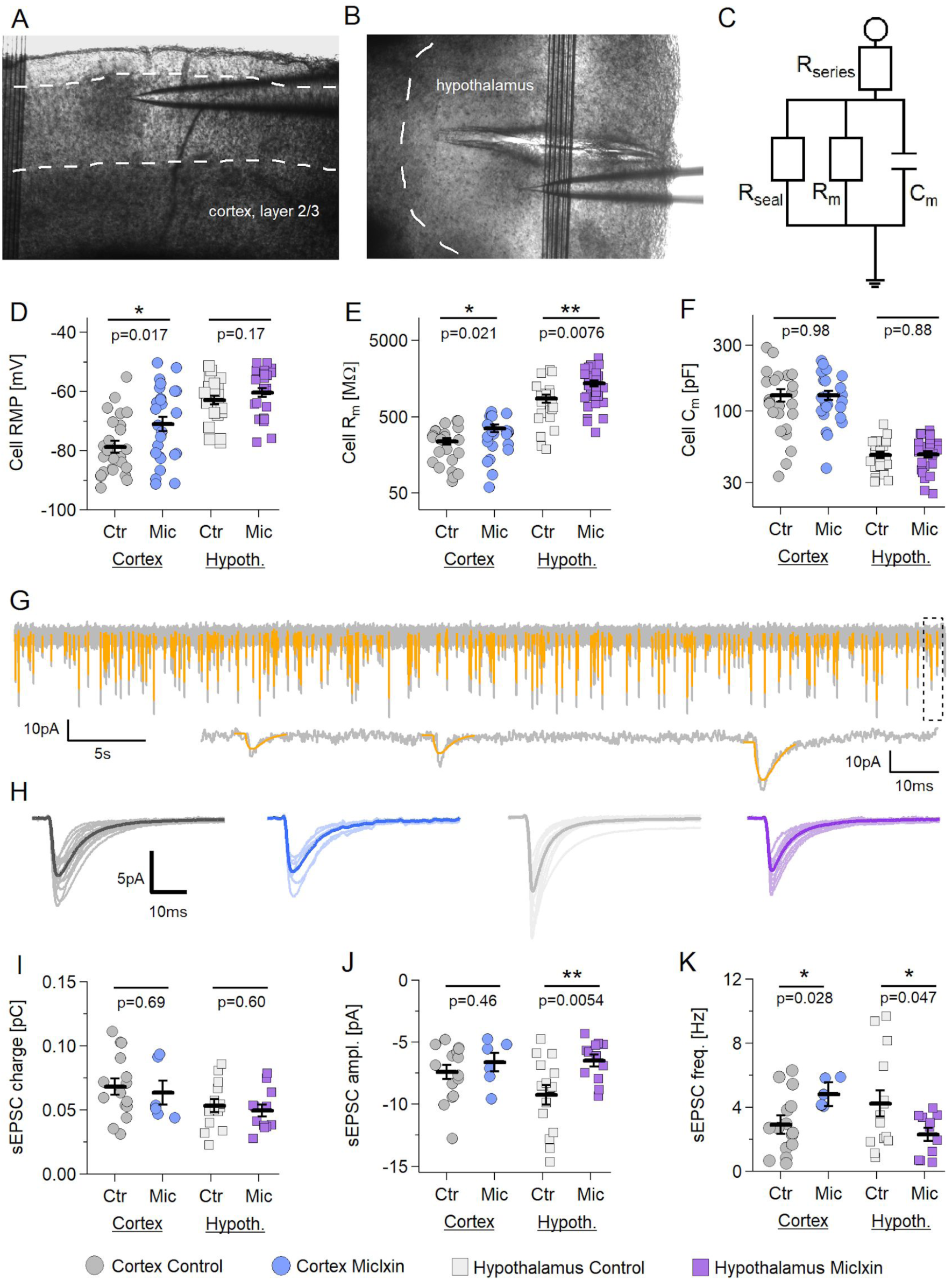
Miclxin alters intrinsic properties of cortical and hypothalamic pyramidal neurons. A) Representative DIC image of the cortical patching area. 4× magnification. B) Representative DIC image of the hypothalamic patching area. 4× magnification. C) Simplified model of major cellular resistances and capacitance. D) Resting membrane potential (RMP) of recorded cortical and hypothalamic pyramidal neurons, compared between control and 50 μM Miclxin groups. Cortex: n = 25 control, n = 25 Miclxin. Hypothalamus: n = 23 control, n = 27 Miclxin. E) Membrane resistance of recorded neurons, as in D. Cortex: n = 24 control, n = 23 Miclxin. Hypothalamus: n = 24 control, n = 29 Miclxin. F) Total capacitance of recorded neurons, as in D. n, is the same as in E). G) Representative 60 s sEPSC recording in a control hypothalamic pyramidal neuron (V_h_ = −85 mV). H) Individual averaged sEPSC waveforms and group means (in bold). I) Total charge transfer for averaged sEPSCs in recorded cortical and hypothalamic pyramidal neurons, compared between control and Miclxin groups. J) sEPSC amplitudes, as in C. K) sEPSC frequency, as in C. In all graphs, bars represent group average ± SEM. Cortex: n = 15; Miclxin: n = 6. Hypothalamus: n = 15 control, n = 10 Miclxin.

#### 2.8.2 Divergent effects of MIC60 inhibition on synaptic transmission in the cortex and hypothalamus

Given that MIC60 inhibition altered neuronal membrane properties and considering the crucial role of mitochondria in supporting synaptic activity (Harris et al., 2012; Lee and Kim, 2015), we next examined the effect of MIC60 inhibition on excitatory synaptic transmission (Fig. 5G–K). In cortical neurons, miclxin treatment did not significantly alter sEPSC charge transfer (68.30 ± 6.36 fC vs. 63.51 ± 9.31 fC; Student’s *t* test, *p* = 0.686) or amplitude (−7.41 ± 0.58 pA vs. −6.63 ± 0.75 pA; Student’s *t* test, *p* = 0.456) but significantly increased the sEPSC frequency (2.93 ± 0.48 Hz vs. 4.82 ± 0.35 Hz, Student’s *t* test, *p* = 0.028). Similarly, in hypothalamic neurons, miclxin treatment did not affect sEPSC charge transfer (53.43 ± 4.89 fC vs. 49.78 ± 4.65 fC; Student’s *t* test, *p* = 0.597), but significantly reduced the sEPSC amplitude (−9.24 ± 0.77 pA vs. −6.47 ± 0.46 pA, Student’s *t* test, *p* = 0.0054) and sEPSC frequency (4.23 ± 0.81 Hz vs. 2.31 ± 0.39 Hz, Welch’s *t* test, *p* = 0.047). Together, these findings indicate that MIC60 inhibition exerts distinct region-dependent effects on excitatory transmission: increasing the synaptic frequency in cortical neurons while reducing the excitatory input in hypothalamic neurons. MIC60 inhibition produces a clear dissociation between synaptic input and spike output that depends on the brain region. In the cortex, miclxin depolarized the RMP (≈ −79 → −71 mV) and increased input resistance (Rm ≈ 239 → 351 MΩ; Fig. 5D–F). Those passive changes would normally favor action potential generation (a smaller current is required to reach the spike threshold) and are consistent with the observed increase in the sEPSC frequency (≈3.6 → 5.6 Hz; Fig. 5G–K), which reflects either an increase in the presynaptic release probability or greater network activity.

#### 2.8.3 MIC60 inhibition results in decreased frequency of spontaneous firing

In the previous section, we established that MIC60 inhibition greatly reduced the ability of both cortical and hypothalamic pyramidal neurons to fire APs upon current injection. In those experiments, we held the cells at ∼–85 mV (V_h_) to prevent premature firing during the injection steps. Moreover, the 0.5 s current injections represent largely nonphysiological inputs. Therefore, we next investigated how MIC60 inhibition affects the spontaneous firing of neurons when they are maintained at their resting membrane potential (RMP). Numerous neuronal populations fire spontaneously in the absence of stimuli (Häusser et al., 2004); thus, we next sought to test how MIC60 inhibition influences this neuronal property.

First, we categorized the neurons into four groups based on their firing rate at the RMP: not firing (N), occasionally firing (<0.1 Hz, O), continuously firing at a low frequency (>0.1 Hz, CL), and continuously firing at a high frequency (>1 Hz, CH; Figure 6A, representative traces). Miclxin treatment markedly altered the neuronal firing patterns in both the cortex and hypothalamus compared with the control treatment.

**Figure 6:**
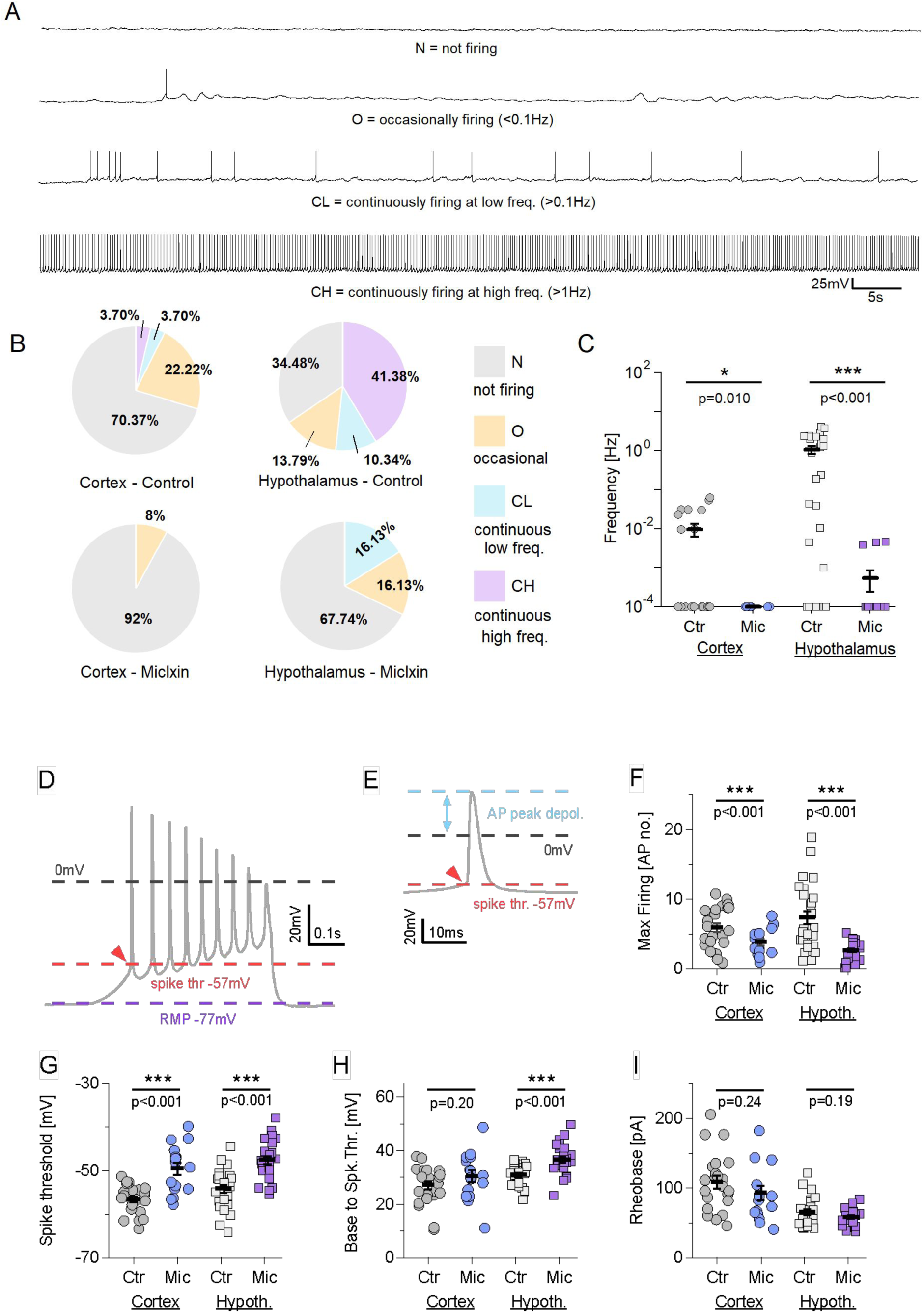
MIC60 inhibition results in decreased frequency of spontaneous firing. A) Four types of spontaneous firing patterns among cortical and hypothalamic neurons, differentiated by firing frequency. B) Percentage of cortical and hypothalamic neurons in each category as in A, compared between control and Miclxin conditions. Cortex: n = 25 control, n = 25 Miclxin. Hypothalamus: n = 31 control, n = 29 Miclxin. C) Scatter plot of firing frequency of cortical and hypothalamic neurons, compared between control and Miclxin conditions. n numbers as in B). In all graphs, bars represent group average ± SEM. *p < 0.05, ***p < 0.001. **Miclxin increases AP spike thresholds and decreases firing rates in cortical and hypothalamic pyramidal neurons.** D) Example of a triangular (ramp) current injection into a cortical pyramidal neuron, to a maximum of 300 pA over 0.5 s. The arrowhead indicates the firing threshold. E) Top: Portion of the trace in D containing the first fired AP. Bottom: Representation of the injected current, with indicated spike threshold and rheobase. F) Comparison of the maximal firing rate triggered by the injection, between control and Miclxin-treated cortical and hypothalamic pyramidal neurons. Cortex: n = 24 control, n = 16 Miclxin. Hypothalamus: n = 26 control, n = 28 Miclxin. G) As in F, for the spike threshold. n is the same as in F). H) As in F, for the difference between the baseline voltage (∼−85 mV) and spike threshold. n is the same as in F). I) As in F, for the rheobase. n is the same as in F). In all graphs, bars represent group average ± SEM.

In controls, cortical pyramidal neurons fired only rarely when recorded at their RMP, whereas hypothalamic principal neurons tended to fire spontaneously much more often (Figure 6), likely because of their more depolarized RMP and higher Rm (see Figure 5D, E). The majority of cortical neurons were nonfiring (N: 70.37%), and those that fired (29.63%; Figure 6B) did so only occasionally. Neurons that were firing continuously at a high frequency (>1 Hz) accounted for only 3.70% of the total. Following miclxin treatment, the nonfiring population increased to 92.00%, while the fraction of firing neurons decreased to 8.00%, with no cells firing continuously at either low or high frequencies (Figure 6B). However, the statistical comparison revealed no significant difference between the two populations (Fisher’s exact test, p = 0.160).

In the control hypothalamus, a majority of the neurons (51.72%) fired continuously at frequencies as high as 3–7 Hz, with a total of 65.52% classified as spiking neurons. Like in the cortex, miclxin reduced the number of firing neurons, but the effect was more pronounced: after treatment, 67.74% of the neurons did not spike, and only 32.26% fired, with none at frequencies >1 Hz (Figure 6B). Here, the difference between the control and miclxin-treated groups was highly significant (Fisher’s exact test, p = 2.24E-04), suggesting that MIC60inhibition has a stronger effect on the hypothalamus than on the cortex.

As the next step, we quantified the reduction in firing by analyzing the average firing frequency. In the cortex, the mean firing frequency decreased from 0.010 ± 0.0036 Hz in control neurons to 0 Hz in miclxin-treated neurons. The only two neurons that fired (0.0040 and 0.0045 Hz) were identified as outliers (Figure 6C). In the hypothalamus, the mean firing frequency decreased from 1.058 ± 0.24 Hz to 0.00054 ± 0.00030 Hz (Figure 6C). In both brain regions, the experimental groups differed significantly (Welch’s t test; cortex: p = 0.010; hypothalamus: p = 1.67E-04).

#### 2.8.4 MIC60 inhibition decreases neuronal firing rates and increases the threshold for action potential generation

Given that MIC60 inhibition increased the sEPSC frequency in the cortex but reduced it in the hypothalamus and affected sEPSC amplitudes differently, the underlying mechanism remains unclear. Such changes could arise either from altered synaptic release properties or from differences in firing at the network level. We next examined how neuronal firing properties were affected to distinguish between these possibilities. Therefore, we utilized current injections to assess neuronal responses, beginning with 300 pA ramp protocols to evaluate the firing rate, spike threshold and rheobase (Figure 6D, E). First, we analyzed the maximal firing rate, which was reduced in both the cortex (5.91 ± 0.62 APs in the control group vs. 3.81 ± 0.46 APs in the miclxin group; Student’s t test, p=0.0095; Figure 6F) and the hypothalamus, where the reduction was even more pronounced (7.33 ± 0.94 APs in the control group vs. 2.63 ± 0.28 APs in the miclxin group; Welch’s t test, p=4.47E-05). Additionally, we observed an approximately 6-mV increase in the spike threshold in both the cortex (-41.02 ± 0.70 mV in the control group vs. -34.06 ± 1.38 mV in the miclxin group; Welch’s t test, p=1.64e-04) and the hypothalamus (-38.60 ± 0.89 mV in the control group vs. -32.04 ± 0.89 mV in the miclxin group; Student’s t test, p=3.49E-06; Figure 6G). Interestingly, a significant increase in the difference between the resting membrane potential (RMP) and spike threshold was observed only in the hypothalamus (30.87 ± 0.80 mV in the control group vs. 36.58 ± 1.12 mV in the miclxin group; Student’s t test, p=1.20e-04) but not in the cortex (27.13 ± 1.49 mV in the control group vs. 30.42 ± 2.18 mV in the miclxin group; Student’s t test, p=0.204; Figure 6H). Finally, no significant differences in rheobase were detected in either region (cortex: 108.72 ± 8.78 pA in the control group vs. 92.82 ± 10.06 pA in the miclxin group; Student’s t test, p=0.242; hypothalamus: 65.16 ± 4.21 pA in the control group vs. 58.68 ± 2.44 pA in the miclxin group; Student’s t test, p=0.189; Figure 6I).

#### 2.8.5 MIC60 inhibition alters the input–output relationship in neuronal firing, decreases AP amplitudes and increases the latency between APs

In the previous section, we established that MIC60 inhibition reduced neuronal excitability by lowering firing rates and increasing spike thresholds in both the cortex and hypothalamus without affecting the rheobase. Figure 7A shows a representative example of 5 current injections into a cortical pyramidal neuron in 100 pA increments over 0.5 s. As the next step, we injected cells with a series of current steps in 25 pA increments (Figure 7B) to assess how neuronal firing scales with increasing input to probe whether the reduction in AP generation is uniform. In the cortex, MIC60 inhibition caused a significant reduction in firing for injections of 100 pA and higher, up to 400 pA, when AP suppression began (Figure 7B; p=0.0058 to p=0.40). In the hypothalamus, a reduction in firing was observed for injections of 50 pA and higher and persisted throughout the AP suppression range (Figure 7B; p=2.81E-05 to p=0.015). Consistent with these and previous findings, both cortical and hypothalamic neurons exhibited a decrease in the maximum firing rate (Figure 7C; cortex: 8.94 ± 0.96 APs in the control group vs. 5.58 ± 0.69 APs in the miclxin group; Student’s t test, p=0.014; hypothalamus: 10.32 ± 1.50 APs in the control group vs. 3.51 ± 0.53 APs in the miclxin group; Welch’s t test, p=1.93E-04) and in the total number of fired APs (Figure 7D; cortex: 101.22 ± 15.12 in the control group vs. 58.90 ± 8.91 in the miclxin group; Welch’s t test, p = 0.025; hypothalamus: 86.63 ± 12.64 in the control group vs. 31.03 ± 3.18 in the miclxin group; Welch’s t test, p=2.36E-04).

**Figure 7:**
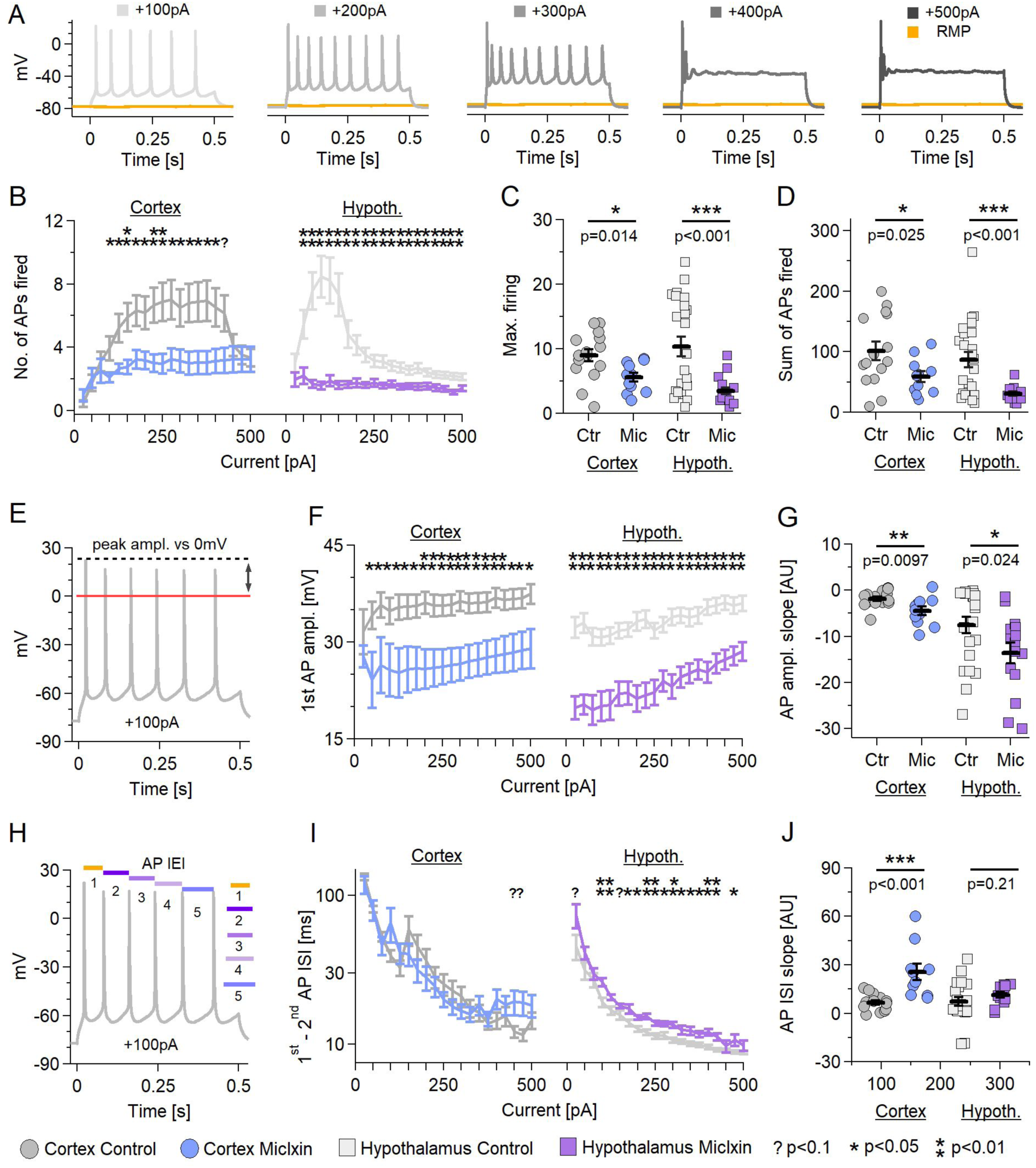
Miclxin reduces neuronal firing, decreases AP amplitudes, and alters AP timing during firing. A) Representative examples of 5 current injections into a cortical pyramidal neuron, in 100 pA increments over 0.5 s. B) Mean input-output curves comparing firing rates of control and Miclxin-treated neurons in cortex and hypothalamus. All injections in +25 pA increments, up to +500 pA. C) Maximum firing rates of neurons in B. D) Total number of APs fired by neurons in B. E) Example of AP peak depolarization measurements during a +100 pA current injection. F) Differences in first AP peak depolarization at each current injection, compared between controls and Miclxin. G) The slope of the linear fit to all AP peak depolarizations, compared between controls and Miclxin. H) Example of time intervals between AP peaks during a +100 pA current injection. I) Summary of differences in time intervals between the 1st and 2nd APs fired at each current injection, compared between controls and Miclxin. J) The slope of the linear fit to all AP time intervals, compared between controls and Miclxin. In all graphs, bars represent group average ± SEM. *p < 0.05, **p < 0.01,< 0.1.

Next, we investigated whether the APs fired had similar amplitudes. An example of AP peak depolarization measurements during a +100 pA current injection is shown in Figure 7E. We found that MIC60 inhibition produced decreased AP amplitudes in both the cortex and hypothalamus (Figure 7F; cortex: p=0.0022 to p=0.047; hypothalamus: p=2.06E-06 to p=3.49E-04) and steeper AP adaptation (Figure 7G; slope; cortex: -1.84 ± 0.48 in the control group vs. -4.53 ± 0.89 in the miclxin group; Student’s t test, p=0.0097; hypothalamus: -7.60 ± 1.76 in the control group vs. -13.65 ± 2.23 in the miclxin group; Student’s t test, p=0.039).

An example of the time intervals between AP peaks during a +100 pA current injection is shown in Figure 7H. Interestingly, when the effects of miclxin on the latency between APs in the cortex were examined, we did not observe significant differences in latency for the first two APs at any current injection (Figure 7I; Student’s t test or Welch’s t test, p=0.083 to p=0.94). However, subsequent APs were progressively more delayed (Figure 7J; 6.49 ± 1.38 ms in the control group vs. 25.53 ± 5.17 ms in the miclxin group; Welch’s t test, p=0.0050). In contrast, in the hypothalamus, the second APs were significantly delayed (Figure 7I; Student’s t test or Welch’s t test, p=8.00E-04 to p=0.045), whereas the latencies of subsequent APs were not significantly different (Figure 7J; 7.14 ± 2.75 ms in the control group vs. 11.32 ± 1.73 ms in the miclxin group; Welch’s t test, p=0.21).

### 2.9 Miclxin alters mitochondrial morphology and dynamics in neurons

We examined mitochondrial morphology and dynamics in iPSC-derived neurons (iNeurons; Figure 8A–H) and Neuro2a (N2A; Figure 8I–N) cells treated with miclxin to determine how miclxin influences mitochondrial organization and function in neurons. High-resolution confocal microscopy images and three-dimensional reconstructions revealed striking alterations in the mitochondrial architecture upon miclxin exposure. In control iNeurons, the mitochondria appeared as elongated and interconnected tubular networks extending along neurites. In contrast, miclxin-treated neurons displayed a predominance of short, rounded mitochondria, indicating extensive fragmentation and a loss of network continuity. The quantitative analysis using Imaris confirmed significant reductions in the mitochondrial area and volume, accompanied by a pronounced increase in sphericity (Figure 8D; ****P < 0.0001; unpaired two-tailed t test), consistent with mitochondrial compaction and morphological remodeling.

**Figure 8:**
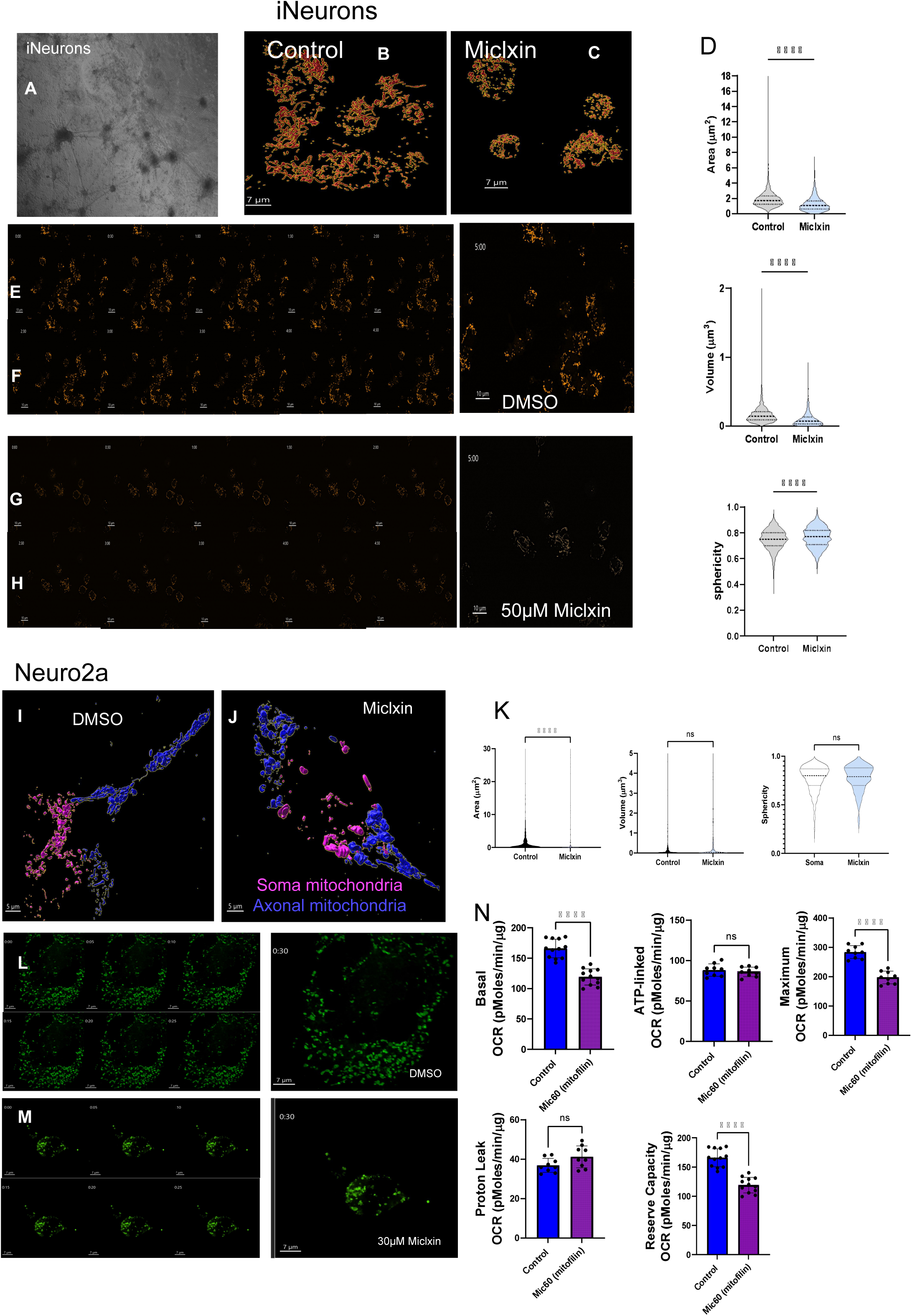
Mitochondrial morphology and dynamics in iPSC-derived neurons (iNeurons) treated with Miclxin. A-C: Representative bright-field and 3D-rendered confocal images of iPSC-derived neurons (iNeurons) under control and Miclxin-treated conditions. Three-dimensional mitochondrial surface reconstructions were generated from Nikon W1 spinning disk confocal images using Imaris to quantify ultrastructural parameters of individual mitochondria. D: Violin plots show mitochondrial *area*, *volume*, and *sphericity* in control and Miclxin-treated neurons (****P < 0.0001, unpaired two-tailed t-test). Miclxin treatment markedly reduced mitochondrial area and volume while increasing sphericity, consistent with mitochondrial fragmentation and compaction. Panels E–H show time-lapse SoRa-mode spinning disk confocal images, capturing mitochondrial dynamics in neurons treated with DMSO (control) or 50 µM Miclxin. Sequential time-stamped frames illustrate loss of elongated mitochondrial structures and reduced motility following Miclxin exposure. Scale bars, 7 µm (top) and 10 µm (bottom). **Mitochondrial morphology and bioenergetic remodeling in Neuro2a (N2A) cells following Miclxin treatment.** I-J: Representative 3D-rendered confocal images of Neuro2a cells treated with DMSO (control) or Miclxin showing distinct mitochondrial distributions within the soma (magenta) and axonal regions (blue). Three-dimensional mitochondrial surface reconstructions and renderings were created in Imaris using data acquired from a Nikon W1 spinning-disk confocal microscope. K: Quantification of mitochondrial *area*, *volume*, and *sphericity* is shown in violin plots (****P < 0.0001, unpaired two-tailed t-test). Miclxin treatment reduced mitochondrial area and volume while increasing sphericity, consistent with mitochondrial fragmentation. Panels below (L,M) show SoRa-mode spinning-disk confocal time-lapse imaging of live N2A cells treated with DMSO or 30 µM Miclxin, demonstrating reduced mitochondrial motility and network continuity over a 30-s interval. N: Lower graphs display Seahorse XF measurements of mitochondrial respiration. Miclxin treatment decreased basal oxygen consumption rate (OCR), ATP-linked respiration, maximal respiration, and spare respiratory capacity, consistent with impaired mitochondrial bioenergetic function. Data are mean ± SEM (*n* = 3–4 biological replicates). Scale bars, 5 µm (top) and 7 µm (bottom).

We performed time-lapse SoRa-mode spinning-disk confocal imaging of live iNeurons to further assess the effect of miclxin on mitochondrial dynamics. In vehicle-treated cells, mitochondria exhibited continuous movement and frequent fusion–fission cycles, which are characteristic of healthy mitochondrial networks. However, within minutes of miclxin exposure, the mitochondria became static and more spherical, and the overall motility along neuronal processes was markedly reduced. These observations indicate that miclxin not only fragments the mitochondrial network but also suppresses its dynamic behavior, suggesting an acute impairment in mitochondrial trafficking and turnover.

A similar trend was observed in N2A cells (Figure 8I–N). Under control conditions, mitochondria were distributed throughout the soma and extended processes, resulting in the formation of elongated networks. Following miclxin treatment, the mitochondria clustered within the soma and were largely absent from the axonal-like projections, reflecting altered intracellular distribution and transport. Quantitative morphometric analyses again showed a reduced mitochondrial area and volume with increased sphericity, reinforcing the morphological signature of fragmentation (Figure 8K). Mitochondrial respiration was assessed using Seahorse XF respirometry to evaluate the functional consequences of these structural changes. Compared with DMSO-treated control cells, N2A cells treated with miclxin exhibited a profound reduction in the basal oxygen consumption rate (OCR), ATP-linked respiration rate, maximal respiratory capacity, and spare respiratory capacity (Figure 8N). These reductions indicate a global decline in oxidative phosphorylation efficiency and bioenergetic reserve.

Collectively, these findings indicate that miclxin strongly disrupted the mitochondrial structure and function in neuronal systems. By inducing mitochondrial fragmentation, immobilization, and bioenergetic suppression, miclxin appears to compromise both the structural integrity and metabolic capacity of neuronal mitochondria, potentially predisposing cells to energy stress and neurodegenerative vulnerability.

## 3. Discussion

In this study, we provide convergent evidence that the mitochondrial contact site and cristae organizing system (MICOS) is a central regulator of neuronal structure and function in the hypothalamus, a brain region critical for systemic homeostasis (Vue 2023; Hinton & Marshall 2024). Using genetic expression analyses, cross-species comparisons, three-dimensional electron microscopy (3DEM), and pharmacological inhibition, we show that MICOS disruption leads to profound ultrastructural remodeling of mitochondria and measurable changes in neuronal excitability (Scudese 2025; Jenkins 2024). These findings integrate molecular, structural, and functional evidence to advance a new framework in which MICOS collapse represents a pivotal step in linking aging, energy imbalance, and susceptibility to neurodegenerative disease (Vue 2023; Hinton 2024). By positioning the hypothalamus at the intersection of mitochondrial biology and systemic physiology, our work extends prior studies of cortical and hippocampal vulnerability in AD and highlights previously unrecognized ancestry- and region-specific differences (Crabtree 2024; Scudese 2025).

## 3. 1 MICOS and AD susceptibility

Our genetic results point to a strong and ancestry-sensitive association between MICOS genes, particularly CHCHD6, and the AD risk (Vue 2023). This result complements previous findings that implicate cortical and hippocampal MICOS deficits in disease, suggesting that subcortical structures may be equally, if not more, vulnerable to MICOS collapse (Vue, 2024; Scudese, 2025). Importantly, ancestry-specific variation in MICOS gene regulation was evident, with differential effects observed in African and European American cohorts (Vue 2023). These findings align with broader disparities in AD incidence and presentation (Hinton 2024), suggesting that mitochondrial remodeling in the hypothalamus may contribute to population-specific risk profiles.

CHCHD6 was the strongest signal in our analyses, which is consistent with its established role in cristae maintenance and MICOS assembly (Vue 2023). These genetic associations reinforce the notion that MICOS genes are not merely structural but also act as disease modifiers. The hypothalamus, with its high metabolic load and continuous activity across the lifespan, may be uniquely susceptible to MICOS perturbations (Hinton & Marshall 2024). These data position MICOS dysfunction as a unifying mechanism connecting genetic ancestry, mitochondrial dynamics, and hypothalamic vulnerability in individuals with AD (Vue 2023).

Our analyses revealed that MICOS genes, particularly CHCHD6, have ancestry-dependent associations with the AD risk (Vue 2023). This finding adds a new dimension to the well-established field of APOE-related genetics (He 2016; Xu 2017). APOE ε4 remains the strongest genetic risk factor for AD across populations, but its penetrance is incomplete and varies by ancestry. For example, APOE ε4 confers a higher risk in European and Asian populations than in African American cohorts, suggesting that ancestry-specific modifiers shape its effects (Xu 2014; Yan 2016). Our results suggest that MICOS collapse may act as one such upstream modifier (Vue 2023).

One possible model is that MICOS dysfunction and APOE risk alleles synergize: APOE-driven lipid and amyloid dysregulation increase cellular stress, whereas MICOS deficits compromise mitochondrial ultrastructure, ATP production, and calcium buffering (Vue 2023; Hinton & Marshall 2024). Together, these alterations accelerate hypothalamic dysfunction, amplifying the systemic metabolic imbalance and increasing susceptibility to neurodegeneration (He 2016; Xu 2017). Importantly, the hypothalamus is a site where APOE is expressed and where lipid–mitochondrial crosstalk is crucial for energy regulation (Hinton 2024). Thus, MICOS collapse could intensify APOE-dependent vulnerability in energy-regulating circuits (Vue 2023).

In addition to APOE, genome-wide association studies (GWASs) have identified additional AD risk loci involved in endosomal trafficking (BIN1 and PICALM), lipid metabolism (CLU and ABCA7), and immune signaling (TREM2) (Crabtree 2024). MICOS dysfunction provides a plausible convergent pathway: impaired mitochondrial–ER tethering and a disrupted cristae architecture could exacerbate vesicular trafficking deficits (BIN1 and PICALM), alter lipid handling (CLU and ABCA7), and increase microglial stress responses (TREM2) (Hinton et al. 2023; Lam et al. 2021; Shao et al. 2024). An ancestry-sensitive MICOS collapse thus represents a systems-level modifier that links diverse risk alleles to a common failure of cellular energetics and neuronal excitability (Vue 2023).

By integrating APOE and other loci into this framework, we propose that MICOS dysfunction is not merely an independent vulnerability but a genetic amplifier that interacts with major AD risk genes (He 2016; Xu 2017). This provides a mechanistic basis for the observed disparities in AD incidence across populations and positions MICOS as a therapeutic target that could mitigate risk even in genetically predisposed individuals (Hinton 2024).

### 3.2 Aging-related MICOS remodeling and evolutionary divergence

Our cross-species comparisons revealed striking differences in the trajectory of the expression of MICOS-related genes with age. In mammals, the expression of MICOS, fusion, and fission genes decreases with age, whereas in flies, the expression of these genes is upregulated (Vue 2023; Crabtree 2024). This divergence may reflect distinct evolutionary strategies for regulating lifespan. In long-lived mammals, the downregulation of these genes may represent a maladaptive collapse under chronic stress, whereas in short-lived species, their upregulation could serve as a compensatory attempt to preserve mitochondrial integrity (Hinton 2024).

The hypothalamus emerges as a particularly vulnerable structure, showing accelerated decreases in MICOS levels compared with cortical regions (Vue 2023; Hinton & Marshall 2024). This pattern is consistent with the role of the hypothalamus in coordinating endocrine and metabolic responses that are energy intensive and highly dependent on mitochondrial efficiency (He 2016; Xu 2017). It also suggests that MICOS collapse may represent a key molecular node linking aging, energy imbalance, and susceptibility to neurodegeneration (Hinton 2024; Scudese 2025).

From an evolutionary perspective, this divergence may reflect different pressures on energy allocation. Flies rely on rapid metabolic shifts to survive, whereas mammals must sustain energy balance over longer periods. The reliance of the mammalian hypothalamus on tightly regulated mitochondrial networks makes it especially sensitive to structural collapse, underscoring why MICOS dysfunction can accelerate systemic aging (Vue 2023; Crabtree 2024).

### 3.2 Ultrastructural remodeling of mitochondria

Our 3DEM analyses provide high-resolution evidence of mitochondrial remodeling in hypothalamic neurons. With MICOS inhibition, mitochondria became smaller, elongated, and more branched, with increased formation of nanotunnels (Jenkins 2024; Scudese 2025). These findings are consistent with recent ultrastructural studies in skeletal muscle, heart, kidney, and brown adipose tissue (Scudese 2025; Vue 2023; Crabtree 2024), which collectively demonstrate that MICOS collapse triggers conserved structural phenotypes across tissues.

The presence of nanotunnels is particularly noteworthy. Traditionally rare in healthy neurons, nanotunnels increased markedly following MICOS inhibition (Jenkins 2024). Previously, researchers have interpreted nanotunnels as intermediates of incomplete fission or stress-induced attempts at intermitochondrial communication (Jenkins 2024). Their emergence here suggests that MICOS collapse drives a pathological remodeling program that interrupts normal fission and may impair organelle quality control (Vue 2023).

These ultrastructural changes have profound functional implications. Smaller and fragmented mitochondria limit ATP production, whereas excessive branching and the presence of nanotunnels reflect instability in network dynamics (Scudese 2025; Crabtree 2024). Given the critical role of the hypothalamus in integrating metabolic and endocrine signals, this remodeling is likely to disrupt systemic homeostasis (He 2016; Xu 2017). Importantly, our data demonstrate that these structural changes occur in neurons, extending previous findings focused on muscle and the heart to the brain (Scudese 2025; Vue 2024).

Our ultrastructural findings in hypothalamic neurons align closely with a growing body of work in nonneural tissues, reinforcing the idea that MICOS collapse elicits a conserved remodeling phenotype across organs. In skeletal muscle, Scudese et al. (2025) reported fragmentation, a reduced cristae density, and excess nanotunnel formation with age, impairing contractile efficiency and recovery. In the kidney, Vue et al. (2023) observed that MICOS deficiency leads to disrupted cristae networks, mitochondrial swelling, and impaired ion transport, ultimately compromising renal homeostasis. Similarly, Crabtree et al. (2024) reported that brown adipose tissue exhibits branching and nanotunnel overproduction when MICOS integrity is lost, diminishing the thermogenic capacity (Cao et al. 2014; Saito, He, Yang, et al. 2016; Xu et al. 2015; Yan et al. 2016; Zhu et al. 2015). Parallel effects have also been described in cardiac tissue, where MICOS dysfunction compromises ATP availability and calcium handling, predisposing patients to arrhythmia (Vue et al., 2023).

Taken together, these reports indicate that MICOS remodeling is a conserved aging signature. Regardless of the tissue type, MICOS collapse manifests as mitochondrial fragmentation, branching, and stress-induced nanotunnels (Scudese 2025; Jenkins 2024). However, the functional consequences are organ specific, reflecting the metabolic demands of each tissue (Crabtree 2024). What distinguishes the hypothalamus is its role as a central integrator of systemic physiology. Unlike the muscle, heart, or kidney—where local deficits impair organ performance—the hypothalamus couples mitochondrial integrity directly to whole-body regulation of feeding, stress, and metabolism (He 2016; Xu 2017). Thus, MICOS collapse in this region is not only a neuronal vulnerability but also a systemic amplifier of aging and neurodegeneration, positioning the hypothalamus as a uniquely sensitive site where conserved remodeling translates into global physiological instability (Vue 2023; Hinton 2024).

### 3.4 Metabolic Insults Alter the Electrophysiological Properties of Neurons

Numerous studies published in recent decades have demonstrated the deleterious effects of metabolic insults on the properties and function of central nervous system (CNS) neurons (Camandola and Mattson, 2017). Hypoxia (Nieber, 1999; Burtscher et al., 2021), food deprivation (Varela et al., 2012; Padamsey et al., 2022), impaired glucose transport (McNay and Pearson-Leary, 2020; Kula et al., 2024), inhibition of oxidative phosphorylation (Hellweg et al., 2003), and insulin resistance (Milstein and Ferris, 2021; Ghasemi et al., 2013; Kleinrdders et al, 2015) are all associated with disruptions in the maintenance of membrane properties, synaptic function, and neuronal firing. However, to date, no investigations have specifically addressed the physiological consequences of disrupting the MICOS complex. In the preceding sections, we delineated the effects of aging and MIC60 knockout on mitochondrial morphology. Here, by employing the recently identified MIC60 inhibitor, miclxin (Ikeda et al., 2020), we examined how the pharmacological inhibition of this essential MICOS component influences neuronal physiology. The re-establishment of the membrane resting potential following synaptic transmission constitutes the most energetically demanding process in neurons, accounting for approximately 43% of total ATP expenditure (Harris et al., 2012). Additionally, disturbances in neuronal metabolism are associated with increased frequencies of postsynaptic currents (PSCs) across various conditions, including food deprivation, glycolytic inhibition, impaired glucose transporter 4 function (a model of insulin resistance), and ischemia. These effects are believed to result primarily from hypoglycemia-induced Ca²⁺ accumulation at presynaptic terminals (Lee and Kim, 2015), and additional alterations in PSC properties have been documented after these metabolic challenges. However, the specific changes observed tend to vary depending on the nature of the impairment (Kula et al., 2024; Padamsey et al., 2022; Lee and Kim, 2015). Collectively, these findings underscore the crucial role of presynaptic glycolysis in sustaining synaptic transmission, as glycolysis typically accounts for approximately one-third of the total energy demand at presynaptic terminals (Lujan et al., 2016). Given the potential severity of MIC60 inhibition suggested by our previous findings, we sought to determine whether miclxin administration alters neuronal properties in the cortex and hypothalamus, two developmentally distinct brain areas (Puelles and Rubenstein, 2003).

Electrophysiological recordings following pharmacological MICOS inhibition (miclxin) revealed the direct consequences of structural collapse on neuronal function. We observed alterations in passive membrane properties, altered excitatory synaptic communication, and a shift in the spike threshold, ultimately lowering neuronal firing rates (He 2016; Xu 2017). These results mirror prior observations in hypothalamic neurons where ion channel modulation regulates excitability and systemic physiology (He 2016; Xu 2017). The sensitivity of the hypothalamus to MICOS disruption underscores its role as a metabolic hub. Previous studies from our group have shown that estrogen, serotonin, and potassium channel signaling tightly regulate hypothalamic excitability and the systemic energy balance (Cao 2014; Hinton 2016., Saito et al. 2016; Xu et al. 2015; Yan et al. 2016; Zhu et al. 2015). MICOS collapse represents a distinct but convergent mechanism of failed excitability: instead of modulating receptor or ion channel activity, it alters the organelle-level foundation required for neuronal firing (Hinton 2024; Neikirk 2023).

This organelle-level vulnerability may explain why hypothalamic dysfunction emerges early in aging and AD. The disruption of excitability in energy-regulating circuits can impair feeding, glucose control, and stress responses, setting the stage for systemic dysregulation (Xu 2014; Yan 2016). Together, our results link the structural collapse of the MICOS to electrophysiological impairments, providing a direct mechanistic bridge between mitochondrial remodeling and altered brain function.

The failed excitability we observed following MICOS disruption resonates with the broader literature on ion channel regulation in hypothalamic neurons. Previous studies have established that the modulation of potassium (K⁺), calcium (Ca²⁺), and hyperpolarization-activated cyclic nucleotide-gated (HCN) channels tightly governs firing rates and systemic physiology (He 2016; Xu 2017). For example, altering K⁺ channel activity can shift membrane resistance and spike thresholds, directly modulating hypothalamic output. Similarly, Ca²⁺ channel dynamics influence excitatory drive, whereas HCN channels regulate rhythmicity and pacemaker-like firing in metabolic circuits (He 2016; Xu 2017).

Our findings suggest that MICOS collapse represents a master upstream regulator of this channel landscape. By fragmenting mitochondria, reducing ATP availability, and destabilizing Ca²⁺ buffering, MICOS dysfunction indirectly shifts the operating range of K⁺, Ca²⁺, and HCN channels (Hinton & Marshall 2024). Instead of modulating a single ion channel subtype, MICOS disruption compromises the bioenergetic and ionic scaffolding that underlies all channel activity. This organelle-level vulnerability explains why hypothalamic excitability decreases in parallel with the process of ultrastructural remodeling. In this sense, MICOS collapse acts not as an isolated defect but as a coordinating failure point, integrating mitochondrial health with electrophysiological integrity (Neikirk 2023; Hinton 2024).

### 3. 5 Integrated structure–function–disease model

This model integrates molecular, structural, and electrophysiological evidence across species and tissues. It builds on prior work linking MICOS dysfunction to skeletal muscle, heart, and kidney pathology (Scudese 2025; Vue 2023; Crabtree 2024) and extends it to the hypothalamus, a previously underexplored site of vulnerability. Importantly, organelle interdependence—particularly mitochondria–ER tethering and calcium dynamics—is central to the translation of ultrastructural remodeling into neuronal dysfunction (Hinton & Marshall 2024; Katti 2023).

Our findings also suggest that MICOS collapse destabilizes mitochondria–ER contact sites (MERCS), which are critical hubs for calcium exchange and metabolic signaling (Hinton & Marshall 2024; Katti 2023). MERCS integrity ensures rapid Ca²⁺ buffering, enabling neurons to maintain firing precision under fluctuating energy demands. When MICOS disassembles, disorganized cristae reduce the mitochondrial calcium uptake capacity, effectively decoupling mitochondria from ER stores. This loss of buffering increases cytosolic Ca²⁺ fluctuations, decreases spike reliability, and destabilizes excitability in the hypothalamic circuits that govern feeding and stress responses (Hinton 2024; Neikirk 2023).

Concurrently, we observed an increased prevalence of nanotunnels, thin protrusions connecting mitochondrial fragments (Jenkins 2024). While previous studies proposed that nanotunnels may act as compensatory conduits to facilitate metabolite or genetic exchange between isolated organelles, their persistence in hypothalamic neurons raises critical questions. Do nanotunnels represent an adaptive attempt to maintain communication in the face of fragmentation or a maladaptive byproduct of incomplete fission and stress signaling? In either case, their presence highlights a profound remodeling of mitochondrial networking strategies (Jenkins 2024; Scudese 2025). Thus, disrupted MERCS and excessive nanotunnel formation reflect two sides of the same pathology—organelle-level instability that undermines neuronal excitability and systemic homeostasis (Hinton & Marshall 2024).

## 4. Conclusions

In summary, our findings establish MICOS as a critical determinant of mitochondrial ultrastructure, neuronal excitability, and hypothalamic vulnerability in aging and disease. Combining genetic, ultrastructural, and electrophysiological approaches, we demonstrate that MICOS collapse represents a unifying mechanism linking ancestry, aging, and neurodegeneration. Our results extend prior work on cortical and muscle mitochondria to the hypothalamus, underscoring its role as a central hub where organelle remodeling is translated into systemic dysfunction. These findings position MICOS not only as a marker of disease progression but also as a promising therapeutic target for preserving brain and body health across the lifespan.

## 5. Methods

### 5.1. MICOS complex GREX association with Alzheimer’s disease in BioVU

The Vanderbilt Institute for Clinical and Translational Research at Vanderbilt University Medical Center curates BioVU, a biobank containing genetic data linked to de-identified electronic health records (EHR). (Dennis et al. 2021) The BioVU program has been previously described. Briefly, this voluntary research program collects leftover blood samples during routine patient care visits at clinics across Tennessee, which are then processed for genotyping or genetic sequencing. BioVU currently houses samples for 344,467 individuals and sample collection is ongoing. (Danciu et al. 2014; Roden et al. 2008)

Genotype data for 94,474 BioVU individuals was generated on the Illumina Multi-Ethnic Genotyping Array (MEGAEX) and quality control procedures included filtering for SNP and individual call rates, sex discrepancies, and excessive heterozygosity. (Dennis et al. 2021) Principal component analysis was used to identify individuals of European and African genetic ancestry, using the 1000 genomes populations as reference. (Price et al. 2006) After imputation using the Michigan Imputation server and the Haplotype Reference Consortium (HRC) reference panel, genotype data underwent additional quality control procedures, including filtering for imputation quality, minor allele frequency, and Hardy-Weinberg Equilibrium.(Das et al. 2016; McCarthy et al. 2016) After the removal of genetically related individuals, we identified 65,363 individuals of European genetic ancestry and 12,313 individuals of African genetic ancestry for analysis. Genetically regulated gene expression (GREX) was calculated in BioVU individuals from models built using the genotype-tissue expression (GTEx) version 8 project data, which includes genotype and matched transcriptome data for 838 donors across 49 distinct tissues. (GTEx Consortium 2020) The best performing GREX models based on the highest r2 values from PrediXcan, UTMOST, and JTI approaches were used to model gene expression for CHCHD3, CHCHD6, and OPA1 from BioVU genotype data.(Gamazon et al. 2015; Hu et al. 2019; Zhou et al. 2020)

To evaluate the correlation between Alzheimer’s disease and MICOS GREX, we extracted Alzheimer’s disease status from the de-identified EHR information in BioVU participants. Specifically, we tested phenotypes or phecodes mapped from ICD9/10 (International Classification of Diseases, 9th and 10th editions) billing codes. The phecode for Alzheimer’s Disease (290.11) includes the following ICD9/10 codes: Alzheimer’s disease (331.0, 331.00, G30), Other Alzheimer’s disease (G30.8), Alzheimer’s disease, unspecified (G30.9), Alzheimer’s disease with late onset (G30.1), Alzheimer’s disease with early onset (G30.0). Information regarding the mapping of phecodes to ICD9/10 codes has been previously described and these mappings are available within the PheWAS package in R (version 0.99.5-2 and 3.6.0, respectively) or online (https://phewascatalog.org/phewas/#home). (Carroll et al. 2014; Wu et al. 2019) We required individuals to have at least two documented instances of the ICD codes on unique dates within the medical record to be considered a case (n_European ancestry = 498, n_African ancestry = 43). Controls were required to have none of ICD codes within the medical record (n_European ancestry = 57,028, n_African ancestry =10,438). Logistic regression analysis was used to examine the association between CHCHD3, CHCHD6, and OPA1 GREX (predictor variable) with Alzheimer’s disease status (outcome variable). Covariates within the regression models included principal components (PC1-10), sex, current age, median age of medical record, and genotype batch in the ancestry stratified analyses. The within-tissue Bonferroni p-value adjustment was calculated by correcting the p-value the number of genes tested (n=3) and the number of phenotypes tested (n=1, adjusted p-value threshold for significance = 0.0167). Nominal significance was considered p<0.05.

### 5. 2 Mouse Care Protocol

Mice with a C57BL/6J genetic background were housed and maintained at 22°C with a 12-hour light/dark cycle and provided unrestricted access to water and standard chow. All animal care followed previous protocols^47^ and was approved by the University of Iowa and University of Rochester Animal Care and Use Committees (IACUCs).

### 5.3 Western Blot Analysis

To obtain protein extracts from male mouse hypothalamus tissue, the brain tissue was homogenized in ice-cold RIPA buffer with protease and phosphatase inhibitors. The lysate was centrifuged at 14,000 x g and 4°C for 15 minutes, and the supernatant was collected.

Quantification was performed using a BCA assay, where BCA Assay Reagents A, B, and C (Thermo Scientific) were added to an albumin standard and cell lysates, with two replicates for each protein of interest, before incubation at 37 °C for 30 minutes. Imaging was performed with Gen5.

Tissue lysates were diluted with RIPA to yield lysates with 2 μg protein/μL lysate, then were diluted further with an equal volume of 2x Laemmli Sampler Buffer (Bio-Rad #1610737) with 5% 2-mercaptoethanol to yield sample solutions with a protein concentration of 1 μg/μL. 15 μL of sample was loaded into 26-well 4–20% Criterion™ TGX Stain-Free™ Protein Gels (Bio-Rad ##5678095) in tris-glycine SDS running buffer (Bio-Rad #1610732). Gels were run at 200V until the dye front reached the bottom of the gel. Gels were equilibrated in tris glycine transfer buffer (KD Medical RGF3395) for 15 minutes, and proteins were transferred to 0.2 μm pore PVDF membranes (Thermo Fisher). Membranes were blocked with 5% bovine serum albumin (BSA)-Tris-buffered saline with Tween-20 and incubated overnight at 4°C with primary antibodies: MIC19 (Protein Tech; diluted 1:2000 with 3% BSA in PBST), OPA1 (Thermo Fisher: diluted 1:1000 with 3% BSA in PBST), and DRP1 (Thermo Fisher: diluted 1:1000 with 3% BSA in PBST). Secondary antibodies were diluted to 1:10000 and incubated with the membrane at room temperature for 1 hour: goat anti-rabbit IgG (H+L) (Invitrogen) and imaged with Gen5. Quantification was performed using ImageJ.

### 5.4 Analysis of Age-Dependent MICOS Gene Expression Using Single-Cell RNA-seq and Pseudobulk Modeling

To investigate age related changes to MICOS related gene expression, murine single-cell RNA-seq data were obtained from the Zhang et al. (2024) Science dataset. Processing was performed in R using Seurat (Stuart et al. 2019; Hao et al. 2021). Samples were subset to the two age classes of interest (“3mo” and “21mo”) and preprocessed using Seurat defaults (log-normalization, identification of variable features, scaling, principal component analysis, construction of the nearest-neighbor graph, and UMAP embedding). A curated list of MICOS/mitochondrial dynamics target genes was used to further subset to features of interest.

To test age effects while accounting for cellular composition and replicate structure, we applied a pseudobulk strategy (edgeR/limma-voom; Robinson et al. 2010; Law et al. 2014; Ritchie et al. 2015). For each object, cells were grouped by cell type and raw counts were summed across cells within each group, yielding a gene × (sample×cell-type) count matrix.

DGE lists were constructed from pseudobulk counts (edgeR), and library sizes were normalized using TMM (calcNormFactors). Mean–variance relationships were modeled with voom (voom), and linear models were fit with limma (lmFit). The design matrix modeled the age effect while adjusting for cell-type identity. Empirical Bayes moderation was applied (eBayes), and differential expression (DE) for age was extracted from the Age_Classold coefficient. Unless otherwise noted in the text, significance was based on Benjamini–Hochberg FDR. Age-stratified target-gene expression was visualized using Seurat’s DotPlot, grouping by Age_Class. To improve visual contrast for highly expressed genes, average expression was capped at 2 before color scaling. Axes labels were replaced with gene symbols, and genes surpassing adj. P < 0.05 in the matched DE results were labeled with an asterisk.

### 5.5 RT-qPCR in *D. melanogaster*

We measured Marf, Drp1, MIC19, and MIC60 transcripts in the brain of laboratory-evolved accelerated aging and control cohorts of *D. melanogaster* at 21 days of age. In *D. melanogaster*, all cDNA samples were diluted to a standard concentration of 1ng/µl before data collection. Three replicates from each population (A and C) were randomly selected after filtering for DNA concentration and purity (260/280 ratio >1.8). Samples were run in duplicate on a CFX96 Touch platform (Bio Rad Laboratories, Hercules, CA). Each well contained 7.5 μL of 2X iTaq SYBR Green Supermix (Bio Rad Laboratories), 3 μL of ddH2O, 0.75 uL of each primer, and 3 μL of template cDNA. Data was analyzed using the ΔΔCt method and normalized to the Drosophila gene rp49/RpL32.

### 5. 6 RT-qPCR in mouse and human samples

Using TRIzol reagent (Invitrogen), total RNA was isolated from tissues and further purified with the rNeasy kit (Qiagen Inc). RNA concentration was determined by measuring absorbance at 260 nm and 280 nm using a NanoDrop 1000 spectrophotometer (NanoDrop products, Wilmington, DE, USA). Approximately 1 μg of RNA was reverse-transcribed using a High-Capacity cDNA Reverse Transcription Kit (Applied Biosciences, Carlsbad CA). Quantitative PCR (qPCR) was then performed using SYBR Green (Life Technologies, Carlsbad, CA)^48^. For qPCR, 50 ng of cDNA was loaded into each well of a 384-well plate, with the reaction carried out on an ABI Prism 7900HT system (Applied Biosystems) with the following cycle: 1 cycle at 95°C for 10 min; 40 cycles of 95°C for 15 s; 59°C for 15 s, 72°C for 30 s, and 78°C for 10 s; 1 cycle of 95°C for 15 s; 1 cycle of 60°C for 15 s; and one cycle of 95°C for 15 s. GAPDH normalization was used to present the data as fold changes.

### 5.7 SBF-SEM

SBF-SEM preparation was performed as described previously (A. Marshall, Damo, et al. 2023; A. G. Marshall et al. 2023). Brain tissue from the amygdala and hypothalamus were removed from mice aged 3 months and 2 years and fixed in 2% glutaraldehyde with 0.1M cacodylate buffer and processed using a heavy metal protocol.

Samples were submerged in 3% potassium ferrocyanide for 1 hour at 4⁰ C, washed with deionized (DI) H2O, and placed in 2% osmium tetroxide for an hour at 4⁰ C. After further washing with DI H2O, samples were immersed in filtered 0.1% thiocarbohydrazide for 20 minutes, washed in DI H2O, and submerged in 2% osmium tetroxide for an additional 30 minutes. After incubation in 1% uranyl acetate at 4 °C overnight, samples were incubated in 0.6% lead aspartate for 30 minutes at 60 °C, then dehydrated in an ethanol-graded series. After fixation, brain samples were infiltrated with epoxy Taab 812 hard resin, immersed in fresh resin, and polymerized at 60 °C for 36-48 hours. Resin blocks were then sectioned for transmission electron microscopy (TEM) to identify regions of interest. Samples were trimmed, attached to an SEM stub, and placed in a FEI/Thermo Scientific Volumescope 2 SEM. Between 300-400 serial thin sections of 0.09 μm per sample block were collected (Garza-Lopez et al. 2021; Hinton et al. 2023; Neikirk and et al. 2023; Lam et al. 2021; A. Marshall, Krystofiak, et al. 2023).

### 5.8 Segmentation and Quantification of 3D SBF-SEM Images with Amira

SBF-SEM images were manually segmented in Amira to perform 3D reconstruction, as previously described. SBF-SEM data from at least three independent experiments were acquired for 3D reconstruction of mitochondria in mice. 300-400 orthoslices for each 3D reconstruction were selected and visualized with Amira software to generate 3D reconstructions and perform quantifications (Figure 2). Mitochondria within axons were manually segmented in Amira software, with ∼100 mitochondria being segmented per ROI. Because more mitochondria were present in the 2-year mouse samples, mitochondria were randomized in the 2-year samples, while all mitochondria in the 3-month samples were reconstructed. Axons were also manually constructed in ROIs to qualitatively determine the orientation of mitochondria within the axons. Quantification of mitochondria was performed with Amira software, which contains built-in parameters.

### 5.9 Structure Quantification and Statistical Analysis

All data are biological replicates. Dots signify individual data points unless specified otherwise. Data from label analyses and manual measurements were statistically evaluated using Student’s t-test or the corresponding non-parametric test using GraphPad Prism (San Diego, California, USA). Graphs are presented as means and black bars indicating the standard error of the mean. Levels of statistical significance p < 0.01, p < 0.001, and p < 0.0001 are denoted as *, **, ***, and ****, respectively.

### 5.10 Quantification of TEM Micrographs and Parameters Using ImageJ

Samples were fixed in a manner to avoid any bias, per established protocols.47 Following preparation, tissue was embedded in 100% Embed 812/Araldite resin with polymerization at 60 °C overnight. After ultrathin sections (90–100 nm) were collected, they were post-stained with lead citrate and imaged (JEOL 1400+ at 80 kV, equipped with a GatanOrius 832 camera). The National Institutes of Health (NIH) *ImageJ* software was used for quantification of TEM images, as described previously.14,48 Student’s t-test was used to test for statistical significance, ** indicates p< 0.01; and *p< 0.05.

### 5.11 Electrophysiology – Slice Preparation

Coronal brain slices containing the hypothalamus and cortical pyramidal cells were used for all electrophysiological recordings. Mice were deeply anesthetized with a mixture of 3% (v/v) isoflurane in air and subsequently decapitated. The brains were removed and sectioned into 300 μm-thick slices using a Leica VT1200S vibratome in ice-cold ACSF composed of (in mM): 230 sucrose, 1 KCl, 0.5 CaCl₂, 10 MgSO₄, 26 NaHCO₃, 1.25 NaH₂PO₄, 0.04 Na-ascorbate, and 10 glucose; osmolarity 310 ± 5 mOsm, pH adjusted to 7.35 ± 0.05 using HCl. The solution was continuously aerated with carbogen (95% O₂ / 5% CO₂) for at least 30 minutes before use.

The slices were then transferred to a BSC-PC slice chamber (Harvard Apparatus, USA) filled with artificial cerebrospinal fluid (ACSF) containing (in mM): 124 NaCl, 2.5 KCl, 2 CaCl₂, 2 MgSO₄, 26 NaHCO₃, 1.25 NaH₂PO₄, 0.04 Na-ascorbate, and 10 glucose; osmolarity 305 ± 5 mOsm/kg, pH adjusted to 7.35 ± 0.05, and aerated with carbogen. The slices were incubated in ACSF prewarmed to 32 °C for 15 minutes, then gradually cooled to room temperature (23 ± 2 °C) over the following 45 minutes.

After recovery, individual slices were placed in another BSC-PC chamber containing either aerated ACSF or ACSF supplemented with 50 µM Miclxin (MedChemExpress, USA). Slices were incubated with Miclxin for at least 60 minutes, with an average incubation time of approximately 105 minutes. After incubation, slices were transferred to a submerged recording chamber mounted on the stage of an upright microscope (SliceScope, Scientifica, UK) equipped with infrared differential interference contrast (IR-DIC) optics. Recordings were conducted at room temperature, with slices continuously superfused with aerated ACSF at a rate of ∼2.5 ml/min using a peristaltic pump.

### 5.12 Whole-Cell Patch-Clamp Recordings

Cortical (layer 2/3) or hypothalamic pyramidal neurons were selected for recording based on their morphology. Their firing patterns and physiological characteristics were compared against reference data available at https://neuroelectro.org/. Neurons exhibiting significant deviations from properties typical of pyramidal neurons in the two regions were excluded from further analysis.

Patch pipettes were fabricated from borosilicate glass capillaries (BF1-86-10, Sutter Instruments, USA) using a vertical puller (PC100, Narishige, Japan). Pipettes were filled with an internal solution containing (in mM): 136 K-gluconate, 4 Na₂ATP, 2 MgCl₂, 0.2 EGTA, 10 HEPES, 4 KCl, 7 phosphocreatine, and 0.3 Na₃GTP; osmolarity 280–290 ± 5 mOsm/kg, pH adjusted to 7.3 ± 0.05 with KOH. Pipette resistance ranged from 4 to 7 MΩ.

During voltage clamp (VC) recordings, cells were held at a membrane potential (V_h_) of −85 mV using a Multiclamp 700B amplifier (Molecular Devices, USA). A liquid junction potential of 15.52 mV was subtracted in all recordings. Pipette capacitance was compensated after seal formation, but cell capacitance and series resistance (R_s_) were not compensated. In current clamp (CC) mode, recordings were bridge-balanced to 50% of the resting potential R_s_. Following cell opening (whole-cell configuration), the cell was held in I = 0 mode to monitor resting membrane potential (RMP) for approximately 180 seconds. The recording was then fitted with a 20-term polynomial in Igor Pro to remove noise and periods of instability and the median value from the fit was reported as the neuron’s final RMP.

To monitor Rs over time, a series of 10 square voltage steps (−10, −5, +5, and +10 mV) was applied every 5 to 10 minutes (V_h_ = −85 mV). Voltage step responses were low-pass filtered at 10 kHz using a Bessel filter and sampled at 20 kHz via an Axon Digidata 1550B digitizer (Molecular Devices, USA). Recordings were included only if they met the following criteria: (1) R_s_ after any recording protocol did not increase by more than 20% from its pre-protocol value; (2) R_s_ never exceeded 30 MΩ; and (3) the offset drift at the end of the session did not exceed ±5 mV.

To determine the spike threshold, each neuron was held at approximately −85 ± 3 mV in current clamp mode and subjected to a 300 pA ramp current injection over 0.5 seconds. The spike threshold was defined as the membrane potential at which the first fully developed action potential (AP) occurred. This protocol was repeated 10 times, and the average threshold was reported for each neuron.

To assess firing properties, neurons were subjected to a series of 0.5-second square current injections (25 pA step increments, up to 500 pA) with a 10-second inter-step interval. The holding current was adjusted as needed to maintain V_h_ at ∼−85 ± 3 mV. Spontaneous excitatory postsynaptic currents (sEPSCs) were recorded in voltage clamp mode at V_h_ = −85 mV during a continuous 5-minute sweep, low-pass filtered at 2 kHz with a Bessel filter and sampled at 10 kHz. All data were acquired using pCLAMP software (Molecular Devices, USA).

### 5.13 Whole-Cell Patch-Clamp – sEPSC Analysis

sEPSC events were automatically detected using a convolution-based algorithm implemented in Fbrain 3.03 (Igor Pro 6, WaveMetrics, USA), with custom macros. The detection template had a rise time constant (τ) of 1.5 ms, a decay τ of 10 ms, and an amplitude of −10 pA. The detection threshold (θ) was set at 4.2× the standard deviation of a Gaussian fitted to the all-point histogram of the convolved trace.

Before analysis, all traces were detrended to eliminate baseline drift and filtered with a digital bandpass filter (0.001–200 Hz) post-convolution. Detected events were visually reviewed, and any not exhibiting typical synaptic kinetics were manually excluded. Cells with a rejection rate exceeding 30% were removed from analysis.

Accepted sEPSC waveforms were extracted as 55-ms segments, consisting of 5-ms pre-onset and 50-ms post-onset windows. Waveforms were baseline-adjusted to the pre-onset period, and peak amplitude and total charge transferred during the first 30 ms were calculated for the averaged sEPSC trace of each cell.

### 5.14 Statistical Analysis – Electrophysiology

Statistical analyses were conducted using GraphPad Prism 10.2.3 (GraphPad Software, USA) or R (R Foundation for Statistical Computing, Austria). Outliers were identified and excluded using ROUT (Q = 5%) or Tukey (IQR * 1.5) methods. Data were tested for normality of residuals and equal variance (homoscedasticity). If both criteria were met, an unpaired t-test was used. If data were not normally distributed but had equal variances, a Mann-Whitney test was applied. If variances were unequal, Welch’s t-test was used. Details on p-values, sample sizes (n for cells, N for animals) are provided in the main text or figure legends. For scatter plots, each point represents an individual cell, with horizontal bars indicating group mean ± SEM. All other graphs present data as mean ± SEM.

### 5.15 Cell Culture and Treatment

Human Embryonic Kidney (HEK293) and Neuro-2a (N2A) cells were cultured under standard conditions. For experimental procedures, cells were treated with one of the following: DMSO (vehicle control), Miclxin (a novel MIC60 inhibitor).

#### iPSC Culture

Engineered doxycycline-inducible neurogenin-2 (NGN2) human induced pluripotent stem cell (iPSC)-derived neurons were utilized in this study. Briefly, iPSCs, maintained on E8 medium (Life technologies #A1517001) underwent a pre-differentiation protocol at passage #15 as follows: iPSCs were dissociated using Accutase (Thermo Fisher Scientific, #A1110501) and 2.0–2.5 × 10⁶ cells per well were plated in N2 medium (Gibco #11330-032) supplemented with Y-27632 (10 μM) (R&D Systems, #1254) and doxycycline (2 μg/mL) (Sigma #D9891-1G) onto 6-well tissue culture plates pre-coated with growth factor-reduced Matrigel (Fisher Scientific, #CB40230A; 1 ml per well of 1 mg diluted in 6 mL DMEM/F-12, Gibco #11330-032). Media was changed daily.

After 3 days of pre-differentiation, cells were dissociated using Accutase and replated at a density of 8.0 × 10⁵ cells per well onto pre-coated 12-well plates in Neuron Medium (1:1 N2:SM1; Gibco #21103049 and StemCell Technologies B27, #05711) supplemented with Y-27632 (10 μM), doxycycline (2 μg/mL), and human BDNF recombinant protein (1 μg/mL; Thermo Fisher Scientific, #PHC7074). Media was changed daily. Neuronal cultures were then maintained for 21 days, with half-medium changes performed on Days 7 and 14. On Day 21, the total medium volume was increased to 2 mL per well, and one-third of the medium was replaced weekly thereafter. At day 28, induced neurons were treated with the MICOS inhibitor Miclxin (30 or 50 μM) or a vehicle control (0.01% DMSO in media) for 15 minutes immediately prior to live-cell imaging to assess complex dynamics.

#### Matrigel coating (D-4)

6-well tissue culture plates were coated with growth factor-reduced Matrigel (Fisher Scientific #CB40230A). Matrigel aliquots (1mg) were thawed and diluted into 6mL of DMEM/F-12(Gibco #11330-032). The Matrigel solution was then distributed at 1mL per well and incubated at 37°C for up to five days.

#### Thawing of iPSCs (D-3)

Cryovials of frozen engineered doxycycline-inducible neurogenin-2 (NGN2) human induced pluripotent stem cells (iPSCs) were thawed were thawed in a 37°C water bath until ∼80% thawed, then 4mL of E8 media (Life technologies #A1517001) supplemented with Y-27632(10uM, R&D Systems, #1254) was added dropwise to the cells. The cells were centrifuged at 1000 rpm for 4 minutes. The supernatant was aspirated, and the cell pellet was resuspended in 6mL of E8 media + Y-27632 (10μM). Resuspended cells were plated onto Matrigel coated plates at a 1:6 split ratio and incubated at 37°C with 5% CO2. After 24h, the medium was replaced with E8 media only, 2mL per well, and replaced every day until cells reached ∼70-80% confluency.

#### Pre-differentiation (D-2 to D-1)

Once cell cultures reached ∼80% confluency, iPSC cultures were prepared for pre-differentiation. The spent medium was manually aspirated from each well and washed with sterile PBS.1mL of Versene(Life Technologies; product #15040) was added to one well, and 1mL of Accutase(Thermo Fisher Scientific, #A1110501) was added to the other five wells followed by incubation at 37°C for ∼3 minutes. Versene was removed and 1mL of E8 media + Y-27632(10uM) was added to gently detach cells. These cells are transferred to 5mL of E8 media + Y-27632(10uM) and plated at a ratio of 1:6 for maintenance. In the remaining five wells, the Accutase was used to gently detach the cells by pipetting over the cell monolayer and then transferred to a conical containing N2 media (Gibco 11330-032, Gibco 17502048) + Y-27632(10uM) + doxycycline (2ug/mL) (Sigma #D9891-1G). The cells were centrifuged at 1000 rpm for 4 min. The supernatant was aspirated, and cells were resuspended in N2 media + Y-27632(10uM) + doxycycline (2ug/mL). Cells were counted via bright light microscopy and seeded at a density of 2–2.5 × 10⁶ cells per well in N2 media + Y-27632(10uM) + doxycycline (2ug/mL). Following seeding, cells were maintained in N2 medium supplemented with doxycycline (2ug/mL) with medium changes for two days. In these two days, medium volumes were doubled (4mL per well).

#### Neuronal Differentiation (D-0)

On D-0, the spent medium was manually aspirated from each well and washed with sterile PBS. PBS was removed and 1mL of Accutase was added to each well, followed by a 3 minute incubation at 37°C. Cells were gently detached by pipetting the Accuatase over the cell monolayer and transferred to 3mL of Neuron Medium (1:1 N2:SM1) (Gibco 21103049, Stem cell technologies B27, #05711)+ Y-27632 (10uM) + doxycycline (2ug/mL) + Human BDNF Recombinant protein (1ug/mL) (Thermo Fisher Scientific #PHC7074). Cells were centrifuged at 1000rpm for 4 min, supernatant was aspirated, and cell pellet was resuspended in Neuronal Media (1:1 N2:SM1) + Y-27632 (10uM) + doxycycline (2ug/mL). Cells were counted and then seeded onto a Matrigel coated 12 well plates at a density of 8.0 × 10⁵ cells per well. Neuronal cultures were maintained for 21 days with half-medium changes performed on Day 7 and Day 14. On Day 21, the total media volume was doubled to 2mL per well, and one-third of the media volume was replaced weekly thereafter until neurons were utilized for experimentation.

### 5.16 Mitochondrial Staining

Following treatment, cells were stained with 150 nM MitoTracker Orange CM-H2TMRos (Thermo Fisher Scientific, #M7511) in pre-warmed media and incubated for 35 minutes at 37°C. After incubation, cells were rinsed with warm media. For morphological analysis, cells were subsequently fixed with a 4% paraformaldehyde solution for 10 minutes at 37°C then rinsed three times with PBS before being mounted.

### 5.17 Microscopy and Image Acquisition

All images were acquired using a Nikon CSU-W1 SoRa spinning disk confocal microscope. This super-resolution system utilizes a microlens-enhanced dual Nipkow disk design to achieve approximately 1.4x higher resolution than standard confocal systems. Images were captured with a 60X objective. For dynamic analysis, live-cell time-lapse imaging was performed over a 1-minute duration to observe mitochondrial fusion and fission events. For 3D morphological analysis, fixed cells were imaged by capturing Z-stacks with a voxel size of 0.209 µm in the X and Y dimensions and 0.352 µm in the Z dimension.

### 5.18 3D Image Analysis

Confocal image stacks were imported into Imaris software (v10.1, Bitplane) for 3D visualization and quantitative analysis. Datasets were verified for correct alignment and scaling before analysis. Mitochondrial networks were reconstructed using the Surface rendering function within the Imaris Measurement Pro module. To segment mitochondrial signals from the background, a machine learning algorithm was used to accurately and consistently identify and threshold the signal across all samples. All 3D reconstructions and quantifications accounted for the anisotropic voxel dimensions. The morphological parameters of mitochondrial volume, surface area, and sphericity were quantified from the generated 3D surfaces.

### 5. 19 Statistical Analysis – Microscopy

Quantitative data were analyzed for statistical significance using the non-parametric Mann-Whitney U test. A p-value of less than 0.05 was considered statistically significant.

## Funding

1. This work was supported by the National Institutes of Health Grant K01NS110981 to N.A.S., and the Department of Defense ARO W911NF2410047 to N.A.S.
2. CZI Science Diversity Leadership grant number 2022-253529 from the Chan Zuckerberg Initiative DAF, an advised fund of Silicon Valley Community Foundation to A.H.J. and O.A.A.; and National Institutes of Health grant HD090061 and the Department of Veterans Affairs Office of Research Award I01 BX005352 to J.G. Howard Hughes Medical Institute Hanna H. Gray Fellows Program Faculty Phase (Grant# GT15655 awarded to M.R.M); and Burroughs Wellcome Fund PDEP Transition to Faculty (Grant# 1022604 awarded to M.R.M). National Institutes of Health Grants: R21DK119879 (to C.R.W.) and R01DK-133698 (to C.R.W.), American Heart Association Grants 16SDG27080009 (to C.R.W.) and 24IVPHA1297559 https://doi.org/10.58275/AHA.24IVPHA1297559.pc.gr.193866
3. The BioVU project at VUMC is supported by numerous sources: including the NIH funded Shared Instrumentation Grant S10OD017985, S10RR025141, and S10OD025092; CTSA grants UL1TR002243, UL1TR000445, and UL1RR024975. Its contents are solely the responsibility of the authors and do not necessarily represent official views of the National Center for Advancing Translational Sciences. Genomic data are also supported by investigator-led projects that include U01HG004798, R01NS032830, RC2GM092618, P50GM115305, U01HG006378, U19HL065962, and R01HD074711, as well as additional funding sources listed at https://victr.vumc.org/biovu-funding/.
4. Genomic data are also supported by investigator-led projects that include U01HG004798, R01NS032830, RC2GM092618, P50GM115305, U01HG006378, U19HL065962, R01HD074711; and additional funding sources listed at https://victr.vumc.org/biovu-funding/. Other funding sources include 2D43TW009744 and R21TW012635 (SKM and AK), the Howard Hughes Medical Institute Hanna H. Gray Fellows Program Faculty Phase (Grant# GT15655 awarded to MRM), and the Burroughs Wellcome Fund PDEP Transition to Faculty (Grant# 1022604 awarded to MRM).
5. This project was funded by the National Institute of Health (NIH) NIDDK T-32, number DK007563, entitled Multidisciplinary Training in Molecular Endocrinology to Z.V.; National Institute of Health (NIH) NIDDK T-32, number DK007563, entitled Multidisciplinary Training in Molecular Endocrinology to A.C.; NSF MCB #2011577I to S.A.M.; The UNCF/Bristol-Myers Squibb E.E. Just Faculty Fund (A.H.J.), Career Award at the Scientific Interface (CASI Award) from Burroughs Wellcome Fund (BWF) ID # 1021868.01 (A.H.J.), BWF Ad-hoc Award (A.H.J.), NIH Small Research Pilot Subaward to 5R25HL106365-12 from the National Institutes of Health PRIDE Program (A.H.J.), Vanderbilt Diabetes and Research Training Center DK020593 (A.H.J.), Alzheimer’s Disease Pilot & Feasibility Program (A.H.J.).
6. ANRF (Anusandhan National Research Foundation), ANRF/ECRG/2024/001042/LS, ANRF/IRG/2024/001777/LS. IISER Tirupati, NFSG. (S.K.M) and by an American Society of Nephrology Kidney Cure Transition to Independence Grant (to C.R.W.).
7. NIH Grants R01HL147818, R03HL155041, and R01HL144941 (A. Kirabo). NIH Grant R00DK120876 (D.T.), Harold S. Geneen Charitable Trust Awards Program (D.T.), Alzheimer’s Association AARG-NTF-23-1144888 (D.T.). NIH Grant R00AG065445 (P.J.), Alzheimer’s Association 24AARG-D-1191292 (P.J.), Wake ADRC REC and Development grant P30AG072947 (P.J.). American Heart Association Grant 23POST1020344 (A.K.). American Heart Association Grant 23CDA1053072 (M. S.).
8. The BioVU project at VUMC is supported by numerous sources, including the NIH-funded Shared Instrumentation Grant S10OD017985 and S10RR025141; CTSA grants UL1TR002243, UL1TR000445, and UL1RR024975 from the National Center for Advancing Translational Sciences. Its contents are solely the responsibility of the authors and do not necessarily represent official views of the National Center for Advancing Translational Sciences. Genomic data are also supported by investigator-led projects that include U01HG004798, R01NS032830, RC2GM092618, P50GM115305, U01HG006378, U19HL065962, R01HD074711; and additional funding sources listed at https://victr.vumc.org/biovu-funding/.23CDA1053072 (M.S.). Its contents are solely the responsibility of the authors and do not necessarily represent the official view of the NIH. The funders had no role in study design, data collection, and analysis, the decision to publish, or the preparation of the manuscript.

## Conflict of interest

The authors declare no conflicts of interest. We would like to acknowledge the Huck Institutes’ Metabolomics Core Facility (RRID: SCR_023864) for use of the OE 240 LCMS and Sergei Koshkin for helpful discussions on sample preparation and analysis. We would also like to acknowledge the Huck Institutes’ Metabolomics Core Facility (RRID: SCR_023864) for use of the OE 240 LCMS and Drs. Imhoi Koo, Ashley Shay, and Sergei Koshkin for helpful discussions on sample preparation and analysis. We would also like to thank UCLA investigators for gifting us old and young human liver samples.

## Acknowledgements

1. would like to acknowledge the Huck Institutes’ Metabolomics Core Facility (RRID: SCR_023864) for use of the OE 240 LCMS and Sergei Koshkin for helpful discussions on sample preparation and analysis. We would also like to acknowledge the Huck Institutes’ Metabolomics Core Facility (RRID: SCR_023864) for use of the OE 240 LCMS and Drs. Imhoi Koo, Ashley Shay, and Sergei Koshkin for helpful discussions on sample preparation and analysis.
2. The INRL would like to thank and recognize the generosity of our donors and their families, as this research is only possible due to their support and willingness to donate. We also wish to thank the National Institutes of Health, the US Department of Defense, the Iowa Neuroscience Institute, the Roy J. Carver Charitable Trust, and the Williams-Cannon Foundation for their generous financial endorsement. The INRL is part of the Department of Pathology, whose support is essential for our operation. **Iowa Neuropathology Resource Laboratory,** Department of Pathology.
3. Confocal microscopy was performed using the Vanderbilt Cell Imaging Shared Resource (supported by NIH grants CA68485, DK58404, and EY08126). The Nikon SoRa Spinning Disk Confocal was acquired through 1S10MH130456-01A1.
4. We thank Dr. Maria Kukley (Achucarro Basque Center for Neuroscience, Bilbao, Spain) and Peter Jonas Lab (IST, Klosterneuburg, Austria) for providing custom macros used in Fbrain 3.03 (Igor Pro 6, WaveMetrics, USA)

## Author Contributions

1. B.S., B.K., and H.L. contributed equally to this work.
2. A.H.J. and N.A.S. conceived the study, designed the experiments, and supervised all aspects of the project.
3. B.S., B.K., H.L., P.V., P.K., A.G.M., S.C., S.T., H.M., M.K., E.S.J., P.M., B.R., N.H.K., C.N., A.R., M.H., L.B., B.T., T.G., J.M.A., S.M.W., L.A.M., S.M., C.W., J.G., R.M., and A.C. performed experiments, data acquisition, and analysis.
4. J.C.S., B.M., J.B., O.K., S.G., E.G.L., C.D., D.-F.D., M.P., and D.H. provided technical expertise, instrumentation, or analytical resources for microscopy, imaging, and data interpretation.
5. A.G.M., B.R., R.M., H.M., J.M.A., and A.H.J. conducted the statistical analysis and prepared the figures.
6. A.H.J., N.A.S., B.S., B.K., H.L., P.V., A.G.M., B.R., and R.M. wrote the manuscript with input from all authors.
7. A.H.J., N.A.S., J.C.S., A.G.M., A.K., M.M.S., C.W., J.G., C.D., and D.-F.D. provided supervision and project oversight.
8. All authors discussed the results, commented on the manuscript, and approved the final version.

## Notes

### Competing Interest Statement

The authors have declared no competing interest.

### Summary of Updates

The acknowledgment section was updated to add and remove grants.

